# Lung cancer-fueled emergency myelopoiesis is characterized by an increase of S100A9^+^ and LCN2^+^ hematopoietic stem and progenitor cells

**DOI:** 10.64898/2026.02.24.707656

**Authors:** Evelyn Calderon-Espinosa, Kirsten De Ridder, Mekhin Carpentier, Kim De Veirman, Daliya Kancheva, Isabelle Scheyltjens, Kiavash Movahedi, Kathleen Van den Eynde, Paul De Leyn, Lieven P Depypere, Sophie Hernot, Yanina Jansen, Cleo Goyvaerts

## Abstract

The pivotal role of tumor infiltrating myeloid cells in lung cancer composition and response to therapy is universally recognized. Nevertheless, their main cradle being the bone marrow (BM), remains vastly understudied owing to the spatiotemporal complexity of hematopoiesis and its hard to access anatomical location. Therefore, the BM niche of lung cancer subjects remains understudied which is why we integrated transcriptional and translational single-cell profiling, ELISA and two-photon microscopy to characterize the medullary hematopoietic compartment in orthotopic lung cancer-bearing mice with validation in human non-small cell lung cancer (NSCLC) samples. In brief we found that lung cancer remotely alters the entire hematopoietic process resulting in higher levels of hematopoietic stem cells (HSCs), myeloid and lymphoid multipotent progenitors (MPPs) and downstream predominance of Granulocyte Monocyte Progenitors (GMP), early Granulocyte Progenitors (GP) and Common Monocyte Progenitors (cMoP) at the expense of mature neutrophils and B cells. Furthermore, a significant increase in the expression and secretion of S100A9 and Lipocalin-2 (LCN2), was characteristic across the entire hematopoietic trajectory in lung cancer-bearing mice and patients. *In vivo* inhibition of S100A9 with Tasquinimod reduced tumor growth, irrespective of its combination with immunotherapy. In addition, it altered the secretion profile of S100A9 but also LCN2 in the BM, suggesting that S100A9 serves as an upstream regulator of LCN2 and holds therapeutic premise to treat immunotherapy refractory lung cancer.

## Introduction

Lung cancer is the leading cause of cancer-related deaths worldwide, with non-small cell lung cancer (NSCLC) accounting for approximately 85% of all cases (Bray et al., 2024; Cui et al., 2025). Despite the profound efficacy and safety advantages of immunotherapy (IT) for advanced NSCLC patients, about 90% present primary or secondary resistance to IT (De Ridder et al., 2025; Zhou & Yang, 2023). Refractory response to IT, including to immune checkpoint inhibitors targeting programmed cell death protein 1/programmed death-ligand 1 (PD-1/PD-L1), has been associated with high infiltration of immunosuppressive myeloid cells (Awad et al., 2018; Calderon-Espinosa et al., 2024; Weber et al., 2018; Zilionis et al., 2019).

As the proliferative and longevity potential of most myeloid cell subsets is low, their ampleness in the lung tumor microenvironment (TME) is mainly warranted via emergency myelopoiesis. The latter entails extra- and intra-medullary activation, expansion, and/or mobilization of hematopoietic stem and progenitor cells (HSPCs), especially the MultiPotent Progenitors 3 (MPP3), alongside the Common Myeloid Progenitors (CMPs) and Granulocyte Monocyte Progenitors (GMPs), which are downstream of the MPP3. This shift towards emergency myelopoiesis is commonly at the expense of the Erythroid-Megakaryocyte-primed MultiPotent Progenitors (EMkMPP) and the Common Lymphoid Progenitors (CLPs) downstream of the MPP4 (Calderon-Espinosa et al., 2024). Hence, in most cancer-related studies, a drastic elevation of cells from the Lin^-^Sca-1^+^c-Kit^+^ (LSK) compartment, monocytes, GMPs and early-stage committed neutrophil progenitors (G0 to G3) have been reported in the bone marrow (BM), blood and even in the TME (Aliazis et al., 2024; Engblom et al., 2017; Giles et al., 2016; LaMarche et al., 2024; Levin et al., 2025; Long et al., 2022; Pu et al., 2016; W. C. Wu et al., 2014; L. Zhao et al., 2018). In line, we and others detected an amplitude of tumor-derived paracrine signals such as G-CSF and MCP-3 that are associated with the instigation of emergency myelopoiesis within the BM (Calderon-Espinosa et al., 2024; De Ridder et al., 2022).

Additional consequences of the fact that lung cancer progression remotely manipulates the BM niche are cancer-related anemia (Long et al., 2022; L. Zhao et al., 2018), alterations of bone metabolism (Engblom et al., 2017), BM-modulated pre-metastatic niche formation, metastasis and resistance to IT (Long et al., 2022). Vice versa, it has been observed that PD-L1 blockade can redefine tumor-instigated emergency myelopoiesis (Boumpas et al., 2024). As such, ample preclinical and clinical studies convey that solid cancer progression is a systemic process whereby a continuous crosstalk between the lung TME and the BM niche fuels the recruitment of an immature and often immunosuppressive myeloid cell population to the remote tumor.

Despite these insights, the molecular mechanisms that characterize tumor-manipulated HSPCs in the BM niche remain poorly understood. In part, this is explained by the difficulty of accessing the BM niche, the low fraction of murine LSK or human CD34^+^ HSPCs in blood and the complexity of mimicking hematopoiesis *in vitro*. This knowledge gap calls for in-depth characterization of the hematopoietic compartment in the BM of lung cancer-bearing subjects and how this spurs the onset of an immunosuppressive milieu. Especially as the tumor-manipulated HSPCs might hold untapped biomarker and target potential. To gain a deeper understanding of this premise, we performed single-cell-resolution profiling through gene expression and spatial analysis of the BM niche in lung cancer-bearing mice and patients.

## Results

### The BM niche of lung cancer-bearing mice shows a significant increase in HSPCs

To appraise which numerical and transcriptional alterations take place within the HSPC population upon lung cancer progression, we conducted hashed single-cell RNA sequencing (scRNA-seq) on hind leg-flushed BM from female orthotopic Lewis lung carcinoma (LLC)-bearing mice and age-matched tumor-free controls (**Extended Data Fig. 1A**). In total, our dataset comprised 6674 and 6867 BM cells from healthy and LLC-bearing mice, resp., which formed at 0.6 resolution Seurat’s anchor-based integration, unsupervised clustering and supervised annotation, 21 clusters corresponding to distinct cell types or stages of differentiation (**Extended Data Fig. 1B**). Based on canonical hallmark genes, clusters were further assigned to one of the following seven hematopoietic cell lineages: hematopoietic progenitor-related (HP), neutrophil-related (Neu), monocyte and dendritic cell-related (MoDC), erythrocyte-related (Ery), basophil and eosinophil-related (BaEo), B cell-related and NK/T cell-related (**Fig. 1A**). When we used Monocle3 to define the trajectories rooting from the HP-related cluster, we corroborated that all other clusters originated from the HP-related cluster (**Fig. 1B**). Main hallmark genes per cluster are depicted in an expression overview for the main canonical marker and 3 additional marker genes per cluster (**Fig. 1C and D**).

**Fig. 1.**
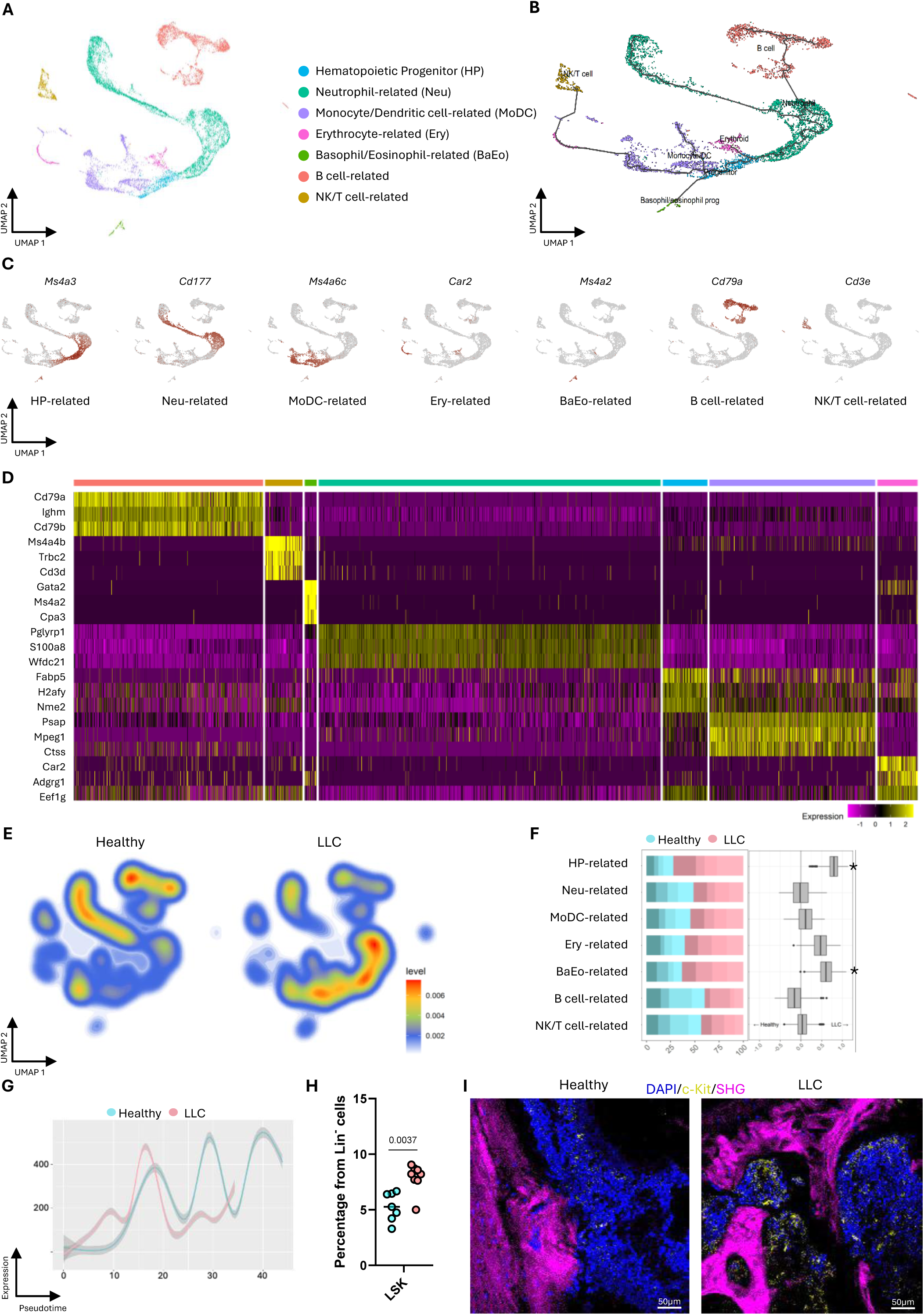
Lewis Lung Carcinoma (LLC) induces a predominance of HSPCs. **(A)** Hashed -seq data from healthy and LLC-bearing female mice (n=4) were processed and integrated using Seurat’s (v.4.3.0) anchor-based integration approach. Harmony-corrected UMAP plots of 6674 and 6867 BM cells resp. show cells clustered at 0.6 resolution. Twenty-one clusters were grouped into 7 cell types according to marker genes. **(B)** Monocle3 trajectory inference of single cells in reduced dimensional space, ordered along pseudotime from the HP-related root and colored by cell type, with the trajectory indicating inferred lineage relationships. **(C)** Expression overview of one marker gene per defined cell type cluster. **(D)** Differential gene expression heatmap showing three additional cluster-specific genes within the cell type. Colors indicate relative expression. **(E)** Density plots showing the cell repartition along the UMAP for each condition. The color gradient represents cell density within the UMAP projection. **(F)** Bar plots depicting cell abundance of each cell type. Four shades of blue and red correspond to the 4 healthy and 4 LLC-bearing mice, resp. By Cacoa package, positive and negative loadings (CoDA loadings per cluster) correspond to over- and under-representation, resp., in LLC-bearing mice compared to healthy mice. Significance levels were determined by a permutation test with Benjamini-Hochberg correction. Asterisk indicates clusters with statistical significance (p <0.05). **(G)** Expression of *Kit* along pseudotime in healthy (blue) and LLC-bearing mice (pink) along the hematopoietic trajectory. Lines show smoothed expression trends, and shaded areas indicate confidence intervals. **(H)** Dot plots depict *ex vivo* flow cytometry abundances of LSK cells from a cohort of gender balanced healthy (n=4F, 3M) and LLC-bearing (n=4F, 4M) mice. Mann-Whitney U test; p-value is shown above. **(I)** Fixed murine humerus slices (30 µm) were stained with DAPI, and c-Kit, and imaged using a two-photon Leica Stellaris microscope at 25X magnification. Bone tissue was visualized via second-harmonic generation (SHG). Scale bar represents 50µm.

Upon evaluation of the density plots along the UMAP projection from each condition, a pronounced shift is observed from predominantly present matured MoDCs, maturing Neu and immature B cells towards HP and myeloid progenitors and reduced B cells in the BM of LLC-bearing mice (**Fig. 1E**). In line, compositional data analysis (CoDA) loadings per cluster confirmed a significant over-representation of the HP-related cluster (**Fig. 1F**). While our scRNA-seq data defined the HP-related cluster by the collective expression of *Ms4a3*, *Fabp5*, *H2afy* and *Nme2,* murine HSPCs are generally defined as LSK cells. Hence, to validate the increased shift of murine HSPCs, *Kit* expression across pseudotime was plotted using Monocle3, confirming increased *Kit* expression in LLC BM cells at early stages of hematopoiesis (**Fig. 1G**). Flow cytometry analysis of the LSK population in flushed BM cells from a cohort of gender balanced healthy and LLC-bearing mice further corroborated the increase in HSPCs (**Fig. 1H**). Finally, fixed murine humerus slices were stained with DAPI and c-Kit to confirm an increase of c-Kit^+^ cells within the BM niche of LLC-bearing mice (**Fig. 1I**). These gene and protein expression data collectively show that the percentage of HSPCs significantly increased in the BM of LLC-bearing mice compared to healthy controls.

### Lung cancer-mediated HSPC abundance rises across myeloid and lymphoid progenitors

Based on Seurat’s unsupervised clustering, the HP-related cluster was defined by the collective and unique expression of *Ms4a3*, *Fabp5, H2afy* and *Nme2*, known for their respective roles in myeloid differentiation (Ishibashi et al., 2018; Jin et al., 2021; Liu et al., 2019), B cell development (Kim et al., 2021) and erythropoiesis (Postel et al., 2009) resp. Hence, this hallmark profile suggests that the HP-related cluster comprises HSPCs for myeloid cells, B cells and erythrocytes. Yet, subcluster analysis of the HP-related cluster showed predominant presence of the MPP3 lineage, comprising CMPs together with their downstream successors, the Myeloid Dendritic cell Progenitors (MDPs) and Monocyte Dendritic cell Progenitors (McDP) (**Extended Data Fig. 1C**). Notably, the other two CMP successors, represented by the GMPs and the MPP4, were not subclustered within the ‘HP-related cluster’ yet assigned to the ‘Neu-related cluster’ and ‘Ery-related cluster’ resp.

As our goal is to understand lung cancer instigated transcriptional changes at the earliest stages of the hematopoietic trajectory, we set out not to limit our evaluation to the HP-related cluster (CMP, MDP and McDP) but include all progenitors across the other six hematopoietic clusters. When we zoomed in on the latter within each subcluster, clear shifts towards higher abundances in the BM of LLC-bearing mice were found for the following progenitors: GMPs and subsequent Neu progenitors G1 and G2 (Neu-related cluster, **Fig. 2A**), MoDC progenitors (MoDC-related cluster, **Fig. 2B**), early pro-B cells (B cell related cluster, **Fig. 2C**) and MPP4 (lymphoid progenitors, **Fig. 2D**). In contrast, the fractions of G3, G4 and resolution-phase macrophages (rM) were significantly reduced, while no clear changes were seen for the EMkMPP nor erythroid progenitors (**Fig. 2A, B, D and E**).

**Fig. 2.**
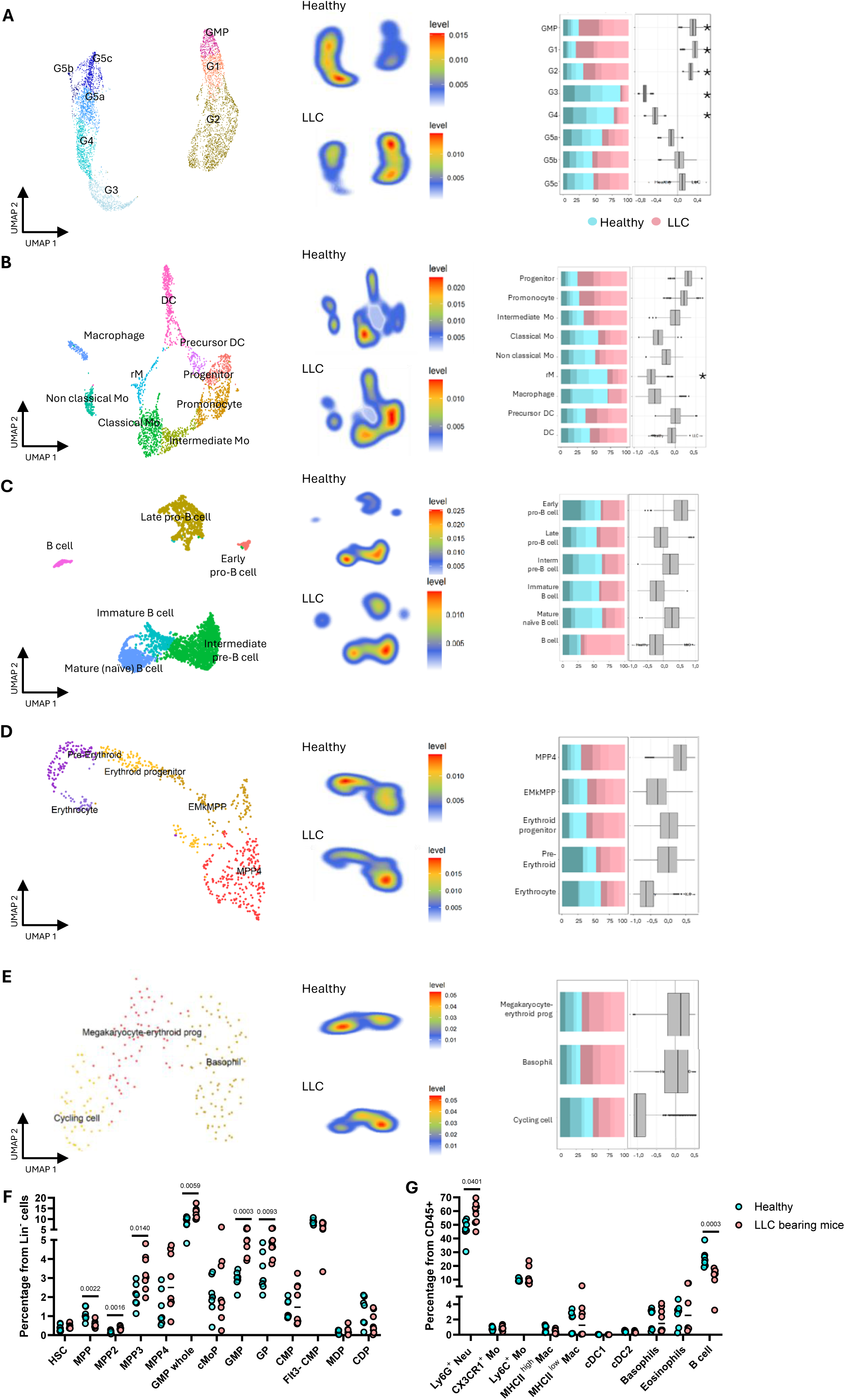
Conservation of progenitor stage across myeloid and non-myeloid cells. (A-E) Left column: Annotations of subclusters when zoomed in on each cell type. Middle column: Density plots showing the cell repartition along the UMAP for each condition, with a color gradient representing cell density within the UMAP projection. Right column: Bar plots depicting the cell abundance of each cell type. Four shades of blue and red correspond to the 4 healthy and 4 LLC-bearing mice, respectively. CoDA loadings per cluster with positive and negative loadings correspond to over- and under-representation resp. in LLC-bearing mice compared to healthy mice. Significance levels were determined by a permutation test with Benjamini-Hochberg correction. Asterisk indicates clusters with statistical significance (p <0,05). Graphs were plotted for **(A)** Neu-related at 0.8 resolution, **(B)** MoDC-related at 0.4 resolution, **(C)** Ery-related at 0.3 resolution, **(D)** BaEo-related at 0.4 resolution, **(E)** B cell-related clusters at 0.3 resolution. Dot plots depict *ex vivo* flow cytometry abundances of **(F)** Lin^-^ progenitor subsets and **(G)** CD45^+^ populations within flushed BM from a cohort of gender balanced healthy (4 females, 3 males) and LLC-bearing (4 females, 4 males) mice. Depicted p-values were obtained upon Mann-Whitney U test.

To validate these alterations in abundance, we optimized a gating strategy for flow cytometry-based analysis of murine HSPC and differentiated immune cells, based on previously defined phenotypic markers (Challen et al., 2021; Liu et al., 2019; Yáñez et al., 2017) **(Extended Data Fig. 2A and B)**. Feature plots were generated to depict the gene expression profile of each flow cytometry marker within the scRNA-seq-defined clusters (**Extended Data Fig. 3** **and 4**). Using BM from a gender balanced cohort of healthy and LLC-bearing mice, we confirmed a significant increase in the abundance of the MPP2, MPP3, GMP, and GP fractions next to an increased trend in MMP4 (**Fig. 2F**). In contrast, the BM of LLC-bearing mice was characterized by only a slight increase in HSCs and even a reduced percentage of MPPs. This suggests that remote LLC presence impacts the MPP reservoir, spurring more differentiation towards MPP2, MPP3 and MPP4. Flow cytometry further confirmed our scRNA-seq findings with increased levels of Ly6C^+^ monocytes (Ly6C^+^ Mo) and Ly6G^+^ neutrophils (Ly6G^+^ Neu) at the expense of B cells (**Fig. 2G, Extended Data Table 1**). Notably, while scRNA-seq analysis revealed that differentiated mature Neu in LLC-bearing mice are reduced, flow cytometry did show a significant increase in Ly6G^+^ Neu. This apparent discrepancy likely reflects the dynamic nature of myelopoiesis in which transcriptional markers of maturation may be low or delayed even in cells that express surface proteins like Ly6G (Grieshaber-Bouyer et al., 2021).

### HSPC abundance rises in the BM of LLC-bearing mice due to impaired differentiation rather than increased proliferation

To better understand why the proportion of HSPCs increases in the BM of lung cancer–bearing mice, we evaluated their expression of the *Mki67* gene, a well-established marker of cell proliferation (Sun & Kaufman, 2018) As we did not find a significant upregulation of this marker in the HP-related cluster, the observed increase in HSPCs is more likely due to a differentiation arrest than enhanced proliferative activity (**Fig. 3A and B**). Notably, in the Neu-related cluster, *Mki67* was significantly upregulated, suggesting that at least for the Neu progenitors proliferation could in part explain their increase.

**Fig. 3.**
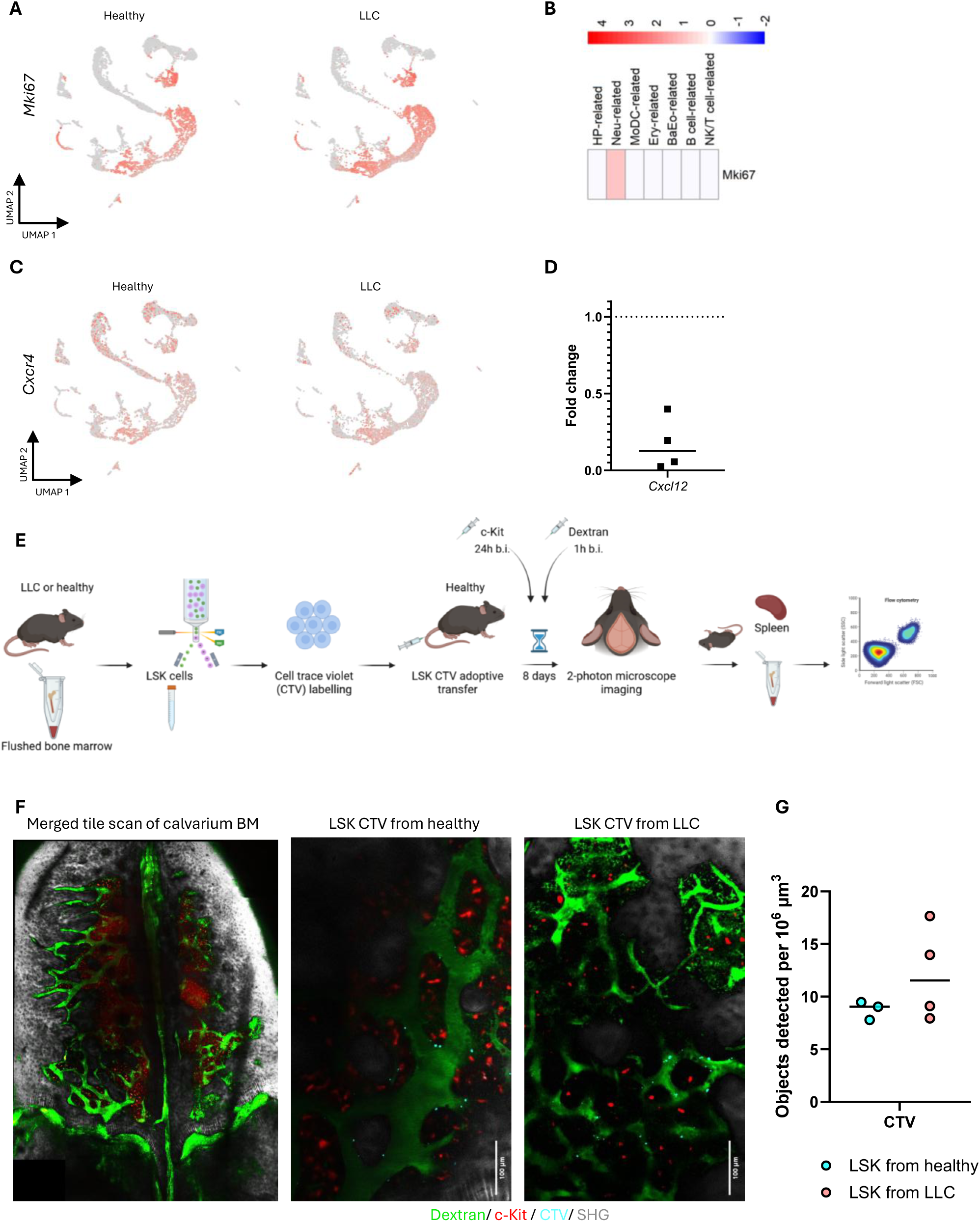
The BM niche, rather than progenitor-intrinsic reprogramming, drives impaired hematopoiesis and progenitor mobilization in lung tumor-bearing mice. **(A)** Feature plots depicting the expression of *Mki67* across individual cells in healthy and LLC-bearing mice. Red color intensity represents the expression level. **(B)** Heatmap of *Mki67* up- or down- regulation expression in the whole dataset. **(C)** Feature plots depicting the expression of *Cxcr4* across individual cells in healthy and LLC-bearing mice. Red color intensity represents the expression level. **(D)** mRNA from crushed BM from healthy (n=2F, 1M) and LLC-bearing mice (n=2F, 2M) was subjected to qPCR for *Cxcl12*. **(E)** Setup of the LSK cell adoptive transfer experiment. C57BL/6 mice were challenged via intrathoracic injection with 5×10^4^ LLC cells. BM from healthy and LLC-bearing mice was collected by flushing hind legs, and 6 × 10^4^ LSK were sorted, labelled with cell trace violet (CTV) and i.v. injected in healthy mice. Eight days after LSK-CTV injection, the calvaria of recipient mice were imaged using a two-photon Leica Stellaris microscope with 25X magnification. Following imaging, mice were euthanized, and the BM and spleen were collected and subjected to flow cytometric analysis to define CTV fractions. **(F)** Left: representative merged tile-scan overview, middle: zoomed-in view of the BM niche from a representative mouse injected with CTV^+^ LSK cells from a healthy donor, right: corresponding zoom-in from a representative mouse receiving CTV^+^ LSK cells from a LLC-bearing donor. One hour prior to imaging, mice were injected with 30mg/kg 500kD dextran-FITC to delineate vasculature and 24hrs prior with 5µg c-Kit-BV711 mAb to show progenitors. Bone tissue was visualized via SHG. Scale bars represent 100µm. **(G)** Images were processed in Fiji using 3D Objects Counter plugin to quantify CTV signal. Statistical significance was assessed using Mann-Whitney U test. Panel **E** created with BioRender.com.

HSPC stemness and immobilization are co-regulated by the stromal BM niche. Therefore we wondered if the differentiation arrest also resulted in a reduced mobilization from the BM niche, as this could explain the observed increase of HSPCs in the BM of LLC-bearing mice. Considering that the stromal BM niche plays a key role in the regulation of HSC mobilization through secretion of HSC retaining factors like CXCL12 (Y. H. Zhao & He, 2025), we wondered if there was any difference in the expression of the CXCL12 receptor CXCR4 on the hematopoietic cells from LLC-bearing mice We found a decrease in the expression of *Cxcr4* across all hematopoietic stages, suggesting reduced sensitivity to CXCL12 and as such increased mobilization (**Fig. 3C**). Using a crushed BM protocol to obtain mRNA from hematopoietic and stromal cells, we further found downregulation of *Cxcl12* in bones from LLC-bearing mice (**Fig. 3D**). These findings suggest that both the hematopoietic cells and stromal BM niche collaborate to stimulate the mobilization of immature progenitors into the bloodstream in LLC-bearing mice, suggesting that the observed rise in HSPCs is due to a differentiation and not mobilization arrest.

To further understand the impact of the BM niche on the HSPCs tendency to remain or egress, we evaluated the BM homing capacity and location of adoptively transferred HSPCs (LSK) from LLC-bearing mice into the tumor-agnostic BM niche of healthy mice (**Fig. 3E**). Specifically, we imaged the calvarium BM of healthy recipient animals (n=3-4), 8 days after they were adoptively transferred with cell trace violet (CTV) labelled LSK cells, sorted from the BM of healthy or LLC-bearing mice. The BM niche was imaged using two-photon microscopy, one day and one hour after mice were injected with anti-c-Kit monoclonal antibody (mAb) and dextran-FITC to visualize stemness and vasculature resp (**Fig. 3F**). Using 3D object counter, we found that the LLC-derived LSKs showed an insignificantly increased tendency to home to the healthy BM niche compared to the healthy animal-derived LSKs (**Fig. 3G)**. Nevertheless, they showed a comparable perivascular location and lack of c-Kit positivity (**Fig. 3F**). This data suggests that LSK cells from LLC-bearing mice do not hold by themselves a progenitor-intrinsic reprogramming that leads to impaired hematopoiesis and progenitor mobilization, but it is the BM niche that governs this impairment. Overall, these data indicate that the stromal component of the BM plays a key role in establishing HSPC stemness maintenance and mobilization in lung cancer.

### *S100a9* and *Lcn2* are significantly upregulated across the entire hematopoietic trajectory

To better understand which transcriptional changes take place within the different BM subclusters of LLC-bearing mice, we examined the differentially expressed genes (DEGs) across the entire BM compartment (**Fig. 4A**). Overall, 190 genes were found to be up- or down-regulated in the BM of LLC-bearing mice compared to healthy controls. Notably, the Cacoa package only defined significant transcriptomic shifts for the different subclusters within the HP- and Neu-related clusters (**Fig. 4B and C**). To define the shared transcriptional changes across all BM clusters, we generated heat maps of the most significantly altered DEGs that were up- or down-regulated in at least two clusters (**Fig. 4D**) and examined their expression patterns within the corresponding subclusters (**Fig. 4E**). *S100a9, S100a8* and *Lcn2* (framed in red on heat maps) were consistently found to be upregulated.

**Fig. 4.**
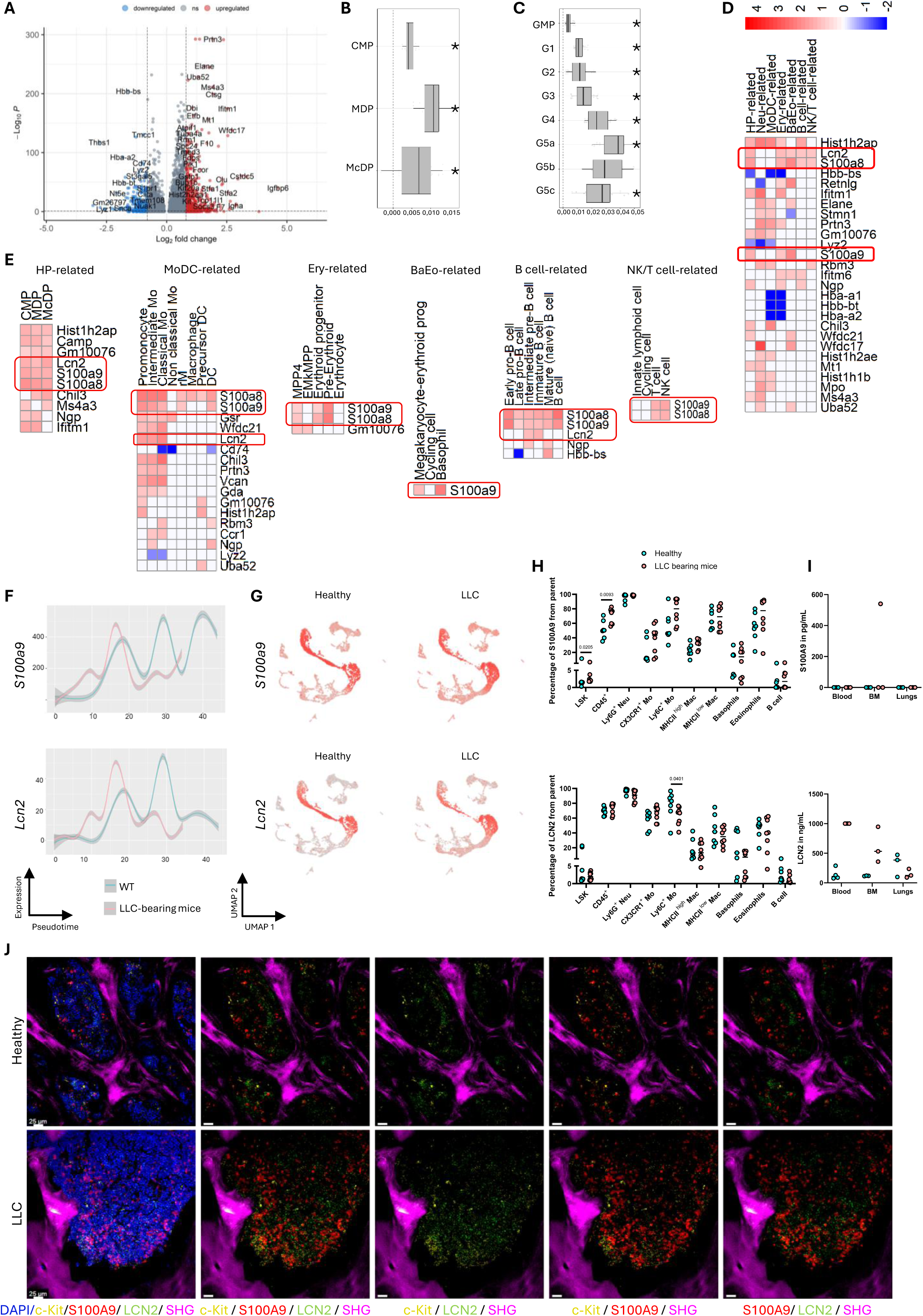
Transcriptional and protein levels of S100A9 and LCN2 are significantly increased in the BM of LLC-bearing mice. **(A)** Volcano plot showing up- and down-regulated genes in the scRNA-seq BM data. Expression shifts between healthy and LLC-bearing mice, as determined by Cacoa package, are shown for the **(B)** HP-related and **(C)** Neu-related subclusters. Significance levels were determined by a permutation test with Benjamini-Hochberg correction. Asterisk indicates clusters with statistical significance (p <0.05). **(D)** Heatmap of ranked up- and down-regulated genes and significant in at least 2 clusters within the entire dataset or **(E)** within specific cell types. **(F)** Plots show expression of *S100a9* and *Lcn2* as a function of pseudotime along the hematopoietic trajectory in the HP-related cluster. Lines show smoothed expression trends, and shaded areas indicate confidence intervals. **(G)** Feature plots depicting the expression of *S100a9* and *Lcn2* across individual cells in healthy and LLC-bearing mice. Red color intensity represents the expression level. **(H)** Bar plots depict *ex vivo* flow cytometry expression of S100A9 and LCN2 from parents, in LSK and immune cell subsets from healthy (n=4F, 3M) and LLC (n=4F, 4M) mice. Multiple Mann-Whitney test comparisons were performed between healthy and LLC-bearing mice per cell type, p-values are indicated on top of comparisons. **(I)** Dot plots showing ELISA-measured concentrations of S100A9 and LCN2 in blood serum, supernatant upon overnight *in vitro* culture of crushed BM and lungs from healthy (n=3) and LLC (n=3) mice. **(J)** Fixed murine humerus slices (30 µm), from healthy and LLC-bearing mice, were stained with DAPI, c-Kit, S100A9 and LCN2, and imaged using a two-photon Leica Stellaris microscope at 25X magnification. Bone tissues were visualized via SHG. Scale bar represents 25µm. For **(A)**, **(D)** and **(E)** adjusted p-value< 0.1 and |Log2FC| > 0.8, red for up- and blue for down-regulated genes in LLC-bearing versus healthy mice.

The genes encoding calcium-binding proteins *S100a9* and *S100a8* were upregulated across the entire hematopoietic trajectory, including all progenitors and differentiated immune cells except for the entire Neu cluster, the non-classical monocytes, EMkMPPs, erythrocytes, innate lymphoid cells and cycling cells. Especially the lack of upregulation of *S100a9* and *S100a8* in the Neu cluster was surprising (**Extended Data Fig. 5A**), as this cluster is characterized by the highest baseline expression levels of *S100a9* and *S100a8* (**Extended Data Fig. 5B**). *Lcn2*, also known as neutrophil gelatinase-associated lipocalin (NGAL), was upregulated in all HP-related cells, promonocytes, intermediate monocytes, classical monocytes, immature and intermediate pre-B cells (**Fig. 4E**). Monocle3-based trajectory analysis of *S100a9* and *Lcn2* expression per cell type and feature plot further confirmed that the highest expression differences between LLC and healthy BM across pseudotime was found within the HP-related cluster and progenitor stages of the entire BM compartment (**Fig. 4F and G**, **Extended Data Fig. 5B and C**).

On the protein level, we were able to confirm that S100A9 is overexpressed in LSK cells and CD45^+^ immune cells in flushed BM of LLC-bearing mice using flow cytometry (**Fig. 4H**). Notably, upon subgating within the immune cell fraction, an increased (insignificant) shift of S100A9 expression was observed in the CX3CR1^+^ monocytes (CX3CR1^+^ Mo) and Ly6C^+^ Mo, in line with the Monocle3 trajectory data (**Extended Data Fig. 5B)**. In contrast, we were unable to show a significant increase in LCN2^+^ cells within the LSK nor CD45^+^ immune cell fraction. Aside from a slight increase within the CX3CR1^+^ Mo, a significant reduction was even found within the Ly6C^+^ Mo next to an (insignificantly) reduced shift of expression in MHCII^low^ macrophages (MHCII^low^ Mac), basophils and B cells (**Fig. 4H**). To further validate secretion of S100A9 and LCN2, ELISAs were performed on murine serum of peripheral blood, supernatants from crushed BM and lung (tumor) samples from healthy and LLC-bearing mice. While secreted S100A9 was only detected in the BM of LLC-bearing mice, secreted LCN2 was detected in all three organs and markedly increased in the blood and BM of LLC-bearing mice, while its levels were reduced in the lungs (**Fig. 4I**). Finally, we confirmed an increased expression of S100A9 and LCN2 via immunohistochemistry on fixed humeri from LLC-bearing mice (**Fig. 4J**). These findings collectively show that S100A9 and LCN2 increase in the BM niche of LLC-bearing mice. This could be shown on the transcriptomic and spatial level for both. Yet via flow cytometry an increase in S100A9 but not LCN2 could be confirmed, while LCN2, but not S100A9, was detectable in sera and BM supernatants. As we simultaneously observe a reduction in the number of LCN2^+^ BM residing cells yet an increase in the level of secreted protein in the supernatant of BM, we hypothesize that LLC stimulates release of LCN2 leading to lower levels of LCN2^+^ cells.

### LLC-mediated upregulation of *Lcn2* and *S100a9* in the BM is *independent of Il17a*

Given the observed involvement of S100A9 and LCN2 in BM alterations during lung cancer progression, and their reported regulation by IL17A (Christmann et al., 2021; J. Xu et al., 2020) we evaluated the expression of these genes by RT-qPCR. Therefore, mRNA was isolated from crushed BM, comprising both hematopoietic and stromal cells (Baccin et al., 2019) from female and male LLC-bearing mice and age-matched healthy controls. In crushed BM from LLC-bearing mice, *S100a9* and *Lcn2* expressions were increased compared to controls, with fold changes ranging from 1.4 to 32, whereas *Il17a* expression showed substantial inter-animal variability (**Fig. 5A and B**). These findings show that the mRNA levels of *Il17a* are not increased in the hematopoietic nor stromal compartment of LLC-bearing mice, partly in line with our scRNA-seq data in which *Il17a* was also not found to be differentially expressed in any of the hematopoietic clusters.

**Fig. 5.**
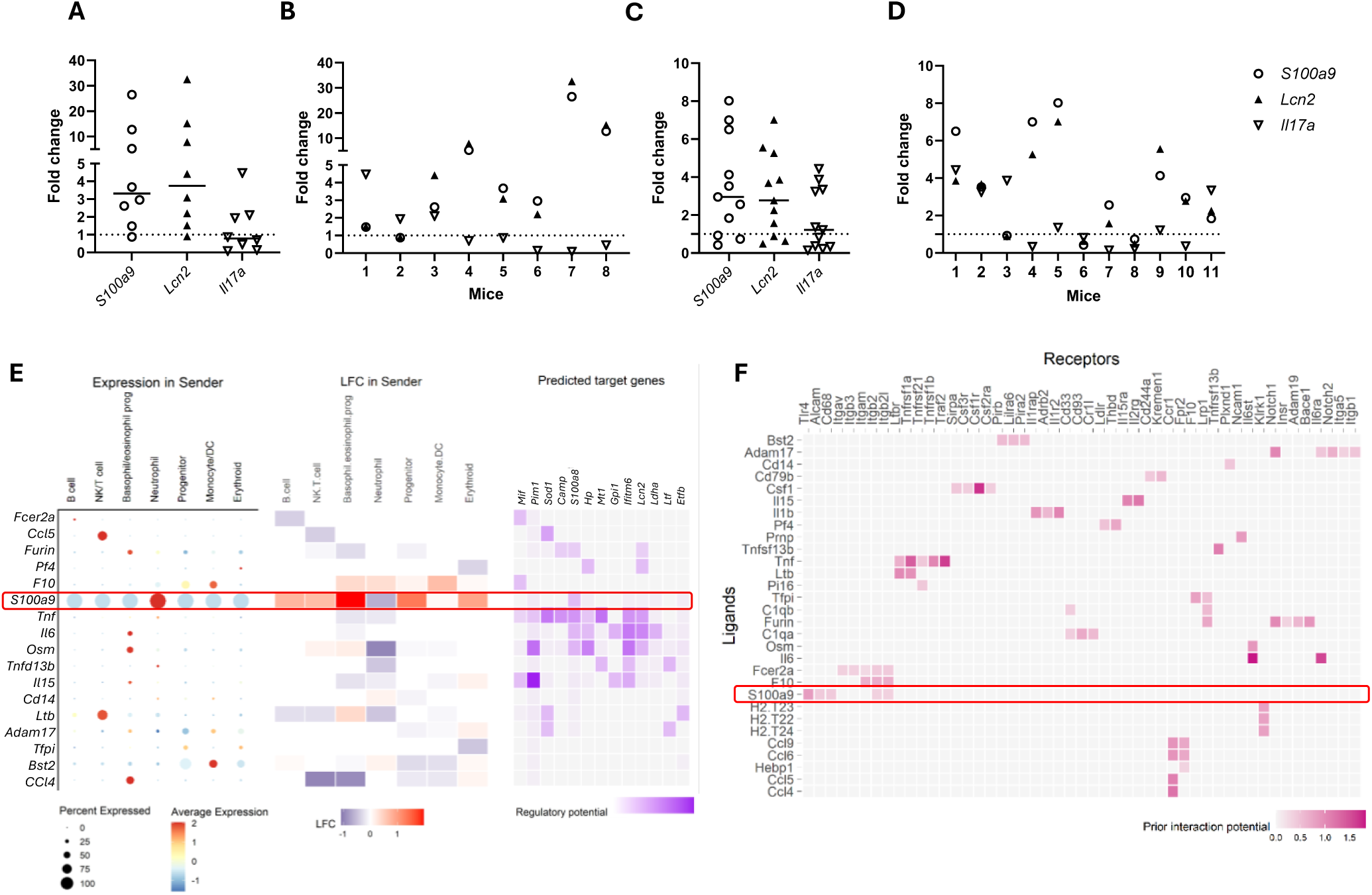
***S100a9* and *Lcn2* are upregulated independently from *Il17a*.** mRNA from **c**rushed BM samples from healthy (n=1F, 2M) and LLC-bearing mice (n=4F, 4M) were used to perform qPCR. (**A)** Summarized and **(B)** individual fold change representation of *S100a9*, *Lcn2* and *Il17a* expression in LLC-bearing mice. Blood from LLC (n=6F, 5M) and healthy (n=3F, 3M) mice were used to perform qPCR. (**C)** Summarized and **(D)** individual fold change representation of *S100a9*, *Lcn2* and *Il17a* expression in LLC-bearing mice. **(E)** NicheNet analysis of the intercellular communication between all cell types (sender cell) and HP-related cell type (receiver cell) based on the scRNA-seq data. Expression of indicated ligands in the sender denotes the regulatory potential between sender-derived ligands and target genes in the progenitor cell type. **(F)** Ligand-receptor matrix shows potential interactions between prioritized ligands and receptors in HP-related cell type.

We next assessed *S100a9*, *Lcn2*, and *Il17a* expression in circulating blood cells. mRNA was isolated from the peripheral blood of female and male LLC-bearing mice and age-matched healthy controls. Consistent with the BM findings, *S100a9* and *Lcn2* were upregulated in LLC-bearing mice; however, the fold change magnitude was more modest than in the BM, with a maximum fold change of 8. In contrast, *Il17a* expression exhibited substantial inter-animal variability, similar to that observed in the BM (**Fig. 5C and D**). Additionally, ELISAs of sera and supernatants from crushed BM and lung samples revealed no detectable IL17A (data not shown).

To understand the relation between the upregulation of *S100a9* and *Lcn2*, we applied NicheNet to evaluate cell-cell interactions within the hematopoietic compartment of the BM of LLC-bearing mice. NicheNet analysis consistently identified *S100a9* as a prioritized ligand and a potential upstream regulator of multiple target genes, including *S100a8*, *Pim1*, and *Lcn2* (**Fig. 5E**), and predicted interactions with several receptors, including *Tlr4*, *Alcam*, *Cd68*, *Itgb2*, and *Itgb2l* (**Fig. 5F**). Although *S100a9* is classically described as being predominantly produced by neutrophils (Sprenkeler et al., 2022), NicheNet analysis suggested that *S100a9* signalling to the HP-related cluster mainly derives from BaEo-related, HP-related and Ery-related populations (**Fig. 5E**). In conclusion *in silico* prediction suggests that S100A9 could act as an upstream regulator of LCN2, making S100A9 the preferred target to evaluate if its inhibition could reduce LLC progression.

### S100A9 inhibitor Tasquinimod reduces lung tumor growth *in vivo*

To further evaluate the effects of S100A9 on hematopoiesis, cancer progression, and immunotherapy, we conducted a gender-balanced *in vivo* experiment. In brief, 7 days after intravenous (i.v.) LLC cell injection, we started treating the mice with anti-PD-L1 mAb or isotype control (IC) intraperitoneally, with or without ad libitum Tasquinimod in drinking water (**Extended Data Fig. 6A**). Tasquinimod is an oral small-molecule immunomodulator primarily investigated for treating castration-resistant prostate cancer and other solid tumors, including ongoing studies in multiple myeloma and myelofibrosis (Fan et al., 2023; Mehta & Armstrong, 2016). Progression of the disease was monitored by micro-computer tomography (micro-CT) (**Fig. 6A**), and body weight (**Extended Data Fig. 6B and C**), twice per week until the humane endpoint was reached. The results did not show a significant difference in survival across the different groups (**Extended Data Fig. 6D**). However, a clear reduction in lung tumor volumes and lung weights could be demonstrated for mice treated with Tasquinimod alone or in combination with anti-PD-L1 mAb (**Fig. 6A-D**). Flow cytometric analysis of cell abundances in BM and spleen did not result in significant alterations. Noteworthy, a trend towards reduced GMP and GP levels in BM in contrast to an increase in Ly6G^+^ Neu could suggest more progenitors shifted towards differentiated cells upon Tasquinimod treatment (**Fig. 6E and F**). In contrast, Tasquinimod seemed to decrease the level of splenic Ly6G^+^ Neu (**Extended Data Fig. 6E and F**).

**Fig. 6.**
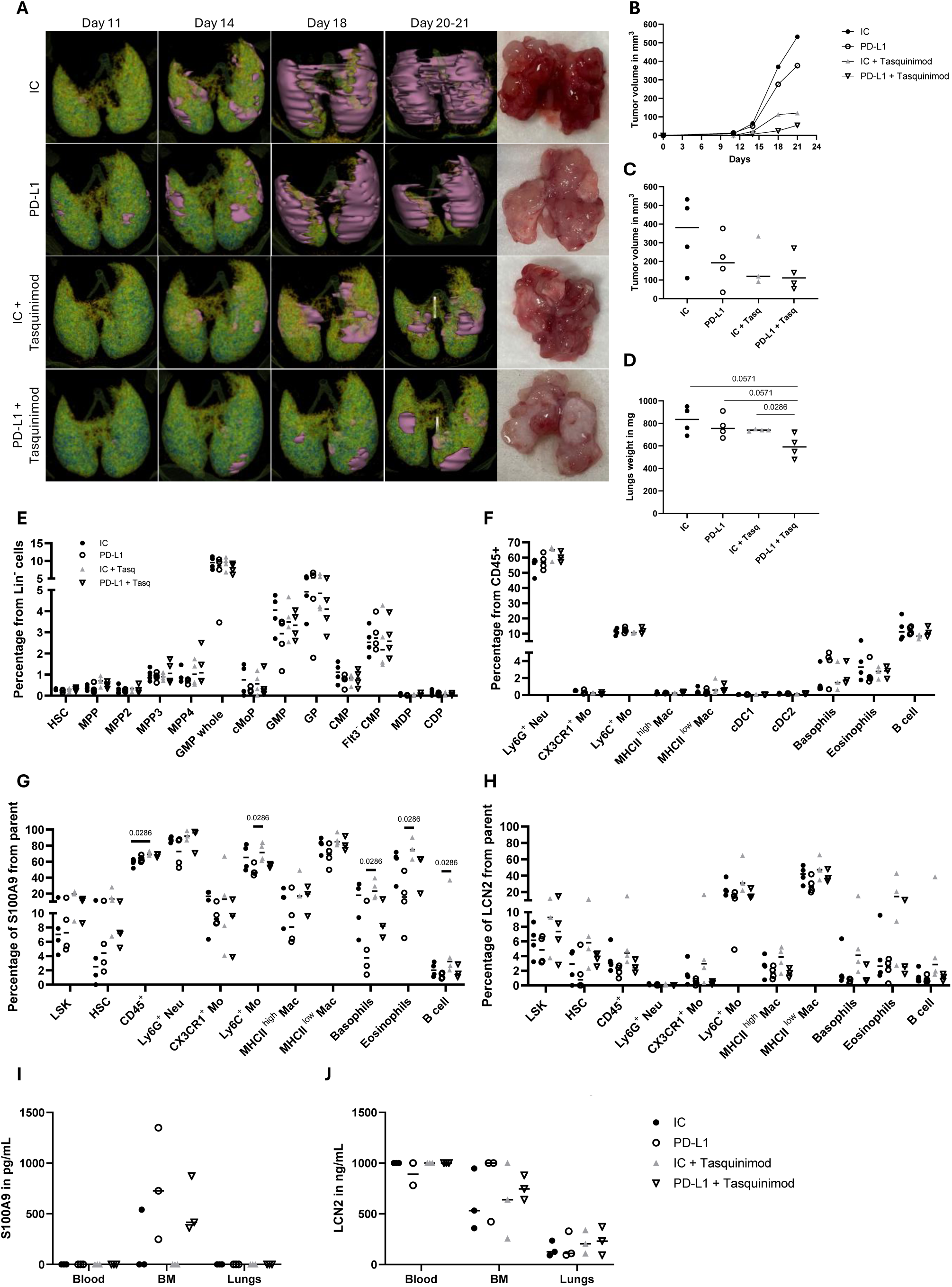
Tasquinimod reduces tumor growth in LLC-bearing mice. LLC-bearing mice were treated with IC (n=2F, 2M), anti-PD-L1 mAb (n=2F, 2M), IC + Tasquinimod (n=2F, 2M) or anti-PD-L1 mAb + Tasquinimod (n=2F, 2M). **(A)** Representative lung micro-CT images acquired during LLC tumor progression from day 11 to day 21 (analyzed with 3D slicer) and corresponding dissected lungs. **(B)** Tumor volumes from day 11 to day 21 in the representative mice of panel A. Tumor volumes calculated via micro-CT (**C**) and lung weights (**D**) of all mice under study at day 21. Dot plots depict *ex vivo* BM flow cytometry abundances of **(E)** Lin^-^ progenitor subsets, **(F)** CD45^+^ differentiated immune cells as well as their **(G)** S100A9 expression and **(H)** LCN2 expression from parent. Depicted p-values were obtained upon Mann-Whitney U test. Dot plots display ELISA-measured concentrations of **(I)** S100A9 and **(J)** LCN2 in blood serum, supernatant upon overnight *in vitro* culture of crushed BM and tumor-bearing lungs.

Next to evaluation of cell abundances, we also scrutinized alterations in S100A9 and LCN2 levels within the BM across treatment groups. Counterintuitively, Tasquinimod monotherapy seemed to increase the levels of S100A9 and LCN2 in the progenitors (LSK and HSC) and differentiated immune cells (**Fig. 6G and H**), suggesting a possibly compensatory mechanism. In addition, a particular pattern was observed: upon anti-PD-L1 mAb treatment as a consistent decrease was found in the percentage of differentiated S100A9^+^ and LCN2^+^ immune cells which could be partly restored when combined with Tasquinimod. Importantly, upon evaluation of secreted S100A9 and LCN2 levels in sera and supernatants from crushed BM and lungs, the opposite pattern was observed. While anti-PD-L1 mAb increased the levels of S100A9 and LCN2 in BM specifically, the addition of Tasquinimod halved this rise (**Fig. 6I and J**). Interestingly, in supernatants from lungs, Tasquinimod did seem to (insignificantly) increase the level of LCN2 protein irrespective of anti-PD-L1 mAb (**Fig. 6J**). As we simultaneously observe a reduction in the number of S100A9^+^ and LCN2^+^ BM residing cells yet an increase in the level of secreted protein in the supernatant of BM, we hypothesize that anti-PD-L1 mAb stimulates release of both proteins, whereas Tasqunimod halts their secretion, leading to higher levels of S100A9^+^ and LCN2^+^ cells. In addition, our data show a tissue-specific response profile to Tasquinimod: whereas the anti-PD-L1 mAb mediated rise in LCN2 secretion is nullified, the opposite is observed in the TME.

### *S100A9*, *LCN2* and *CXCL12* are altered in NSCLC patients

Finally, we validated the alterations of *S100A9* and *LCN2* genes in FFPE rib fragments from operable NSCLC patients. By RT-qPCR, we corroborated that *S100A9* and *LCN2* were altered in NSCLC patients (n=14) compared to healthy controls (n=4). We confirmed that 79% of patients showed upregulation of *S100A9* and 57% showed upregulation of *LCN2* (**Fig. 7A**). Notably, unlike our findings in murine BM samples, these alterations were uncorrelated and patient-dependent (**Fig. 7B**). To assess spatial protein expression, rib fragments from NSCLC and healthy donors were subjected to immunohistochemical analysis. Similarly to the murine results, we observed an increased abundance of CD34^+^ HSPCs in NSCLC patients. Furthermore, higher expression levels of S100A9 and LCN2 were observed in NSCLC patients when compared to healthy donors (**Fig. 7C**). As mRNA was extracted from whole bone samples, comprising hematopoietic and stromal cells, we also studied the expression of the stromal HSC-retention factor CXCL12. In line with the murine data, *CXCL12* was consistently downregulated, indicating their BM niches are also made more prone to increase HSPCs mobilization in NSCLC patients (**Fig. 7D)**.

**Fig. 7.**
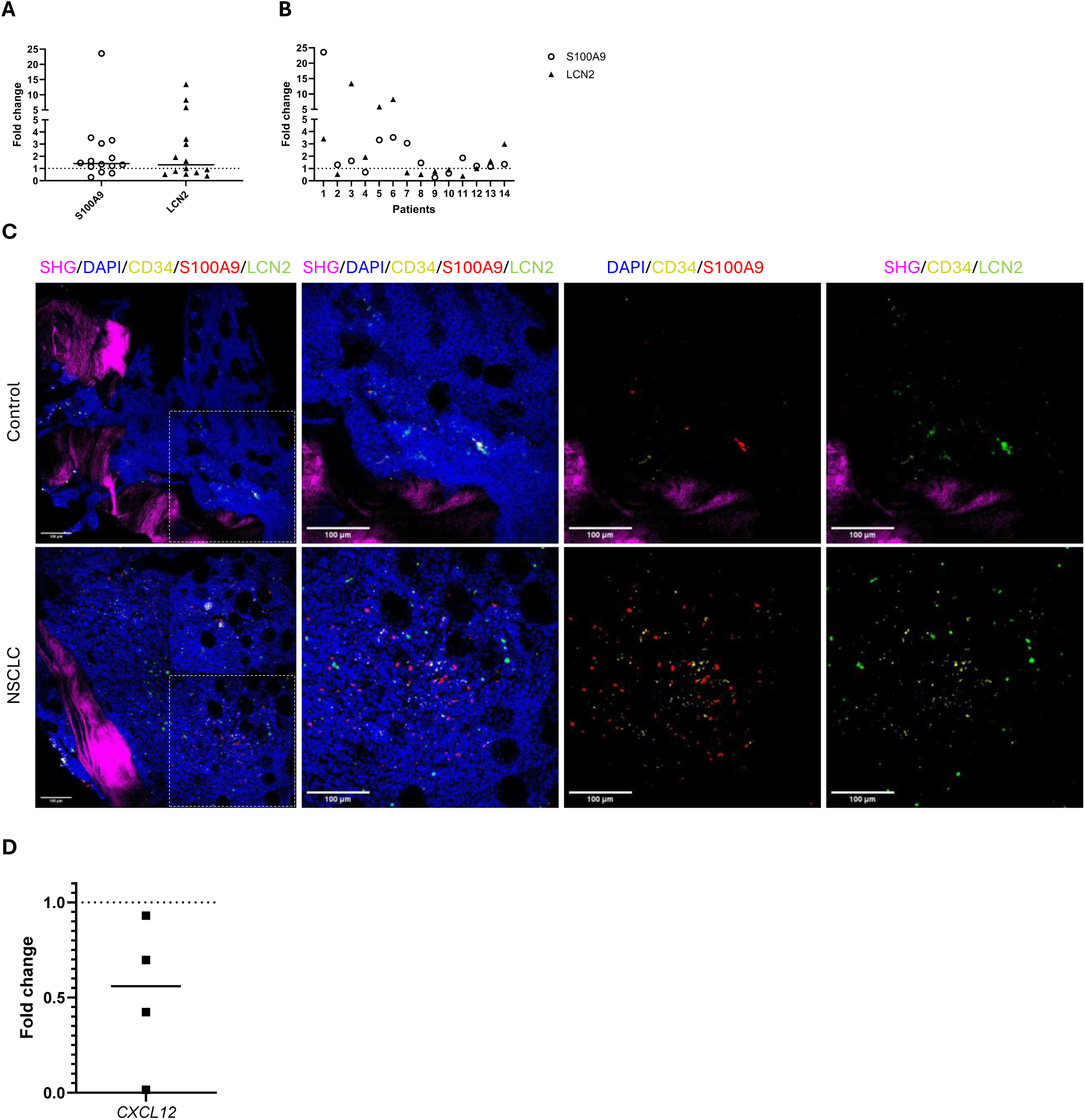
Validation of murine BM alterations during lung cancer progression in NSCLC patients. mRNA from rib fragments from NSCLC patients (n=14) and healthy controls (n=4) were used to perform qPCR. **(A)** Summarized fold change representation of *S100A9* and *LCN2* expression in NSCLC patients. **(B)** Individual expression per patient to define correlations. **(C)** Tile scan overview (left) and zoom in (right) images of fixed rib fragment slices (25 µm) from one representative healthy control and NSCLC patient, stained for DAPI, CD34, S100A9 and LCN2 visualization. Fragments were imaged using a two-photon Leica Stellaris microscope at 25X magnification. Bone tissue was visualized via SHG. Scale bar represents 100µm. **(D)** *CXCL12* qPCR from rib fragments corresponding to healthy (n=3) and NSCLC (n=4) donors.

## Discussion

In this study we integrated single-cell profiling, RT-qPCR, flow cytometry and two-photon microscopy of human and murine BM samples to characterize the transcriptomic and proteomic alterations in the medullary hematopoietic compartment elicited by orthotopic lung tumors. In brief we found that lung cancer remotely alters the entire hematopoietic process resulting in higher levels of HSCs, myeloid and lymphoid MPPs and downstream predominance of GMPs, early GPs and promonocytes at the expense of mature B cells. Furthermore, the expression and secretion of two proteins, S100A9 and LCN2 were characteristic for the entire hematopoietic niche of lung cancer-bearing subjects. *In vivo* inhibition of S100A9 with Tasquinimod reduced tumor growth, irrespective of its combination with immunotherapy. In addition, it altered the secretion profile of S100A9 but also LCN2 in the BM, suggesting that S100A9 serves as an upstream regulator of LCN2 and holds therapeutic premise to treat immunotherapy refractory lung cancer.

Our scRNA-seq findings of the BM suggest that the proportion of HSPCs increases at the expense of differentiated progeny and that this is the result of a differentiation arrest in the HSPCs. Our finding that the proliferation marker *MKi67* was not increased yet the levels of the HSC retention factors CXCR4 and CXCL12 were, further suggests that lung cancer does not increase the proliferation rate of HSPCs but does elevate their stemness maintenance and mobilization into the bloodstream. Although not validated by us, this aligns with the ample studies that reported on increased levels of circulating and tumor-infiltrating immature neutrophils often termed as CD11b^+^Gr1^+^ myeloid derived suppressor cells (MDSC) (Calderon-Espinosa et al., 2024).

Another finding worth mentioning relates to the granulocytes, a collective term for all polymorphonuclear neutrophils, basophils and eosinophils. In line with other studies (Drissen et al., 2016, 2019), Monocle3 based trajectory analysis of our scRNAseq showed that basophils and eosinophils hold a different ontogeny than neutrophils, as they share a developmental origin with the erythroid and megakaryocyte lineages instead of neutrophils. Moreover, it should be noted that our scRNAseq data defined a significant increase in the BaEo-related progenitor clusters in LLC-bearing mice, reflecting the potential understudied importance of these immune cell subsets in lung cancer progression (LaMarche et al., 2024; Zhang et al., 2025).

Upon unbiased evaluation of significant differentially expressed genes on single cell level, *S100a9* was consistently upregulated in the HP-, MoDC-, Ery-, BaEo-, B cell- and NK/T cell-related cluster. This was further validated in bulk murine and human BM via qPCR, on bulk murine blood via qPCR as well as on protein level in the entire LSK and CD45^+^ differentiated immune cell fraction in flow cytometry and murine and human BM samples. A surprising observation was the lack of alterations in the Neu-related cluster as S100A9, often in complex with S100A8 (as calprotectin), is considered a highly specific, and early biomarker for neutrophil activation, comprising up to 45% of neutrophil cytosolic proteins (Sprenkeler et al., 2022). As S100A9 is also known as a myeloid cell-derived pro-inflammatory alarmin, our data suggest that remote tumor growth fuels the pro-inflammatory status of the entire BM niche, including the HSCs and lymphoid compartment holding the NK/T and B cells. Importantly, tumor-specific S100A9 expression has already been associated with poor prognosis and lack of response to immunotherapy (Cheng et al., 2008; Demir et al., 2024; Sinha et al., 2008). The fact that we were only able to detect secreted protein in the supernatant of the BM (not blood nor lung TME), suggests LLC-induced upregulation of BM S100A9 might mainly act in a paracrine fashion.

Next to upregulated *S100a9*, *Lcn2* was also significantly increased in the HP-, MoDC- and B cell-related cluster. Although less pronounced as for *S100A9*, especially for the human samples, this was further validated in bulk murine and human BM via qPCR, in bulk murine blood via qPCR as well as in the serum and BM supernatants of LLC-bearing mice. Flow cytometry did not seem to be sensitive enough for LCN2 detection, or the lack of cellular protein detection was linked to the fact that they already secreted most of their LCN2 making them low in LCN2 but the serum and BM supernatant high in LCN2. The marked increase in LCN2 might in part explain the increase in HSPCs, as it was previously shown that LCN2 increases levels of CXCL12 and SCF (Costa et al., 2017). In addition, LCN2 has been linked to tumorigenesis and is recognized as a potential biomarker across various cancer types (Živalj et al., 2023). Moreover, serum LCN2 levels were correlated with cachexia progression in lung cancer patients (D. Wang et al., 2023). Indeed, we also observed cachexia at late stage of disease in the LLC-bearing mice, in line with results of a meta-analysis in which up to 39% of NSCLC patients have cachexia, and its presence is associated with worse survival (Zhang et al., 2024). Noteworthy though, the level of secreted LCN2 in lungs without tumor was higher than with tumor, warranting further investigation in the impact of increased LCN2 levels in blood and BM niche specifically of lung cancer bearing subjects.

As the simultaneous upregulation of S100A9 and LCN2 has not been described in the BM niche, we used *in silico* prediction of its correlation. While KEGG pathway analysis defined IL17A as a potential upstream regulator of their expression, we did not detect increased IL17A expression in the BM or blood, nor did we detect IL17A in serum or in BM supernatants (data not shown). *In silico*, *S100a9* did in addition act as a potential upstream regulator of *Lcn2*. Akin it was recently shown that S100A9 leads to the activation of the TLR4-p38/MAPK-LCN2 signaling pathway, as such triggering an inflammatory response that contributed to calcium oxalate kidney stone development (Q. Wang et al., 2024).

Considering previous associations of tumor-specific S100A9 with poor prognosis and lack of response to immunotherapy and its potential to act as an upstream regulator of another pro-tumoral protein; LCN2, we set out to investigate the therapeutic potential of Tasquinimod monotherapy or in combination with anti-PD-L1 mAb for LLC-bearing mice. Although Tasquinimod treatment did not improve survival in LLC-bearing mice, a reduction in tumor volume was observed in Tasquinimod monotherapy with or without immunotherapy.

Noteworthy, we opted for an experimental design where Tasquinimod treatment was initiated 7 days after tumor challenge in an aggressive and hard to treat orthotopic LLC lung cancer model. This contrasts preclinical studies where Tasquinimod is provided before tumor onset (Fan et al., 2023). Interestingly, Demir et al., showed that inhibition of extracellular S100A9-mediated TLR4 signaling via Paquinimod even increased tumor growth and reduced anti-PD-L1 mAb efficacy, primarily by decreasing intratumoral Ly6C^+^ Mo without affecting the Ly6G^+^ Neu (Demir et al., 2024). These findings suggest that S100A9-targeted therapies are likely to impact Ly6G^-^ myeloid population within the tumor.

Despite undetectable concentrations of S100A9 in blood serum and TME of LLC-bearing mice, our data showed decreased concentrations of secreted S100A9 in the supernatant of crushed BM when mice were treated with Tasquinimod. In line with these observations, Chung et al., showed that viral nanoparticle vaccines against S100A9 reduce lung tumor seeding and metastasis by reducing S100A9 concentration within the lungs and sera while increasing expression of immunostimulatory cytokines with antitumor function, and the correlation with reduced MDSC populations within the lungs (Chung et al., 2023). Additionally, these findings are in accordance with a trend towards reduced GMP and GP levels in BM in contrast to an increase in Ly6G^+^ Neu that could suggest more progenitors shifted towards differentiated cells upon Tasquinimod treatment, meaning reduced immature suppressor cells. Similarly, Shen et al., showed that in prostate cancer and melanoma, Tasquinimod in combination with immunotherapies provided enhanced response to immunotherapy while reducing MDSC (Shen et al., 2015).

Our data indicate that anti-PD-L1 decreased the number of S100A9^+^ cells in the crushed BM. These results could be partially explained by the observations of Cheng et al., who suggested a functional interplay between S100A9-producing MDSC and the PD-1/PD-L1 axis in myelodysplastic syndromes (Cheng et al., 2019). However, our data also showed that LLC-bearing mice treated with anti-PD-L1 increases the secretion of S100A9 in the BM. The latter observation could explain better that reduced intracellular staining of S100A9 observed by flow cytometry is because of increased secreted protein observed by ELISA.

Previous studies confirmed that S100A9 upregulated the expression of LCN2 in HK-2 cells and that S100A9 deficiency downregulated the renal expression of LCN2 in nephrocalcinosis mice (Q. Wang et al., 2024). Our study shows a similar impact of Tasquinimod on S100A9 and LCN2 expression and secretion in LLC-bearing mice, reaffirming the hypothesis that S100A9 regulates LCN2.

Upregulation of *S100A9* and *LCN2* in human samples from NSCLC patients was not as pronounced as the alterations observed in mice. However, it is important to consider the heterogeneous character of the NSCLC population compared to the syngeneic C57Bl/6J LLC-bearing mice. Moreover, human samples were collected from operable patients at stages I-II of the disease, while mouse samples were collected from early to late stages of the disease.

Although it was not within the scope of this study to investigate which tumor-derived molecular cues install hematopoietic alterations, as ample studies already identified crucial cues such as GM-CSF, G-CSF, M-CSF, MCP-3, SCF, IL-6, IL-8, VEGF, PlGF and IL-1. Downstream of these cues, a prevalent role has been assigned to the CXCL-12-CXCR4 HSC retention pathway within the BM niche where CAR stromal cells hold a pivotal role in the regulation of emergency myelopoiesis (Baccin et al., 2019; Calderon-Espinosa et al., 2024; Mitroulis et al., 2020). Importantly, we observed downregulation of *CXCL12* in mouse and human lung cancer subjects accompanied by downregulation of *Cxcr4* in the BM of LLC-bearing mice. With these findings, we suggest that impaired hematopoiesis in the BM is due to BM niche alteration via lung cancer signaling through the stromal compartment. Therefore, we believe it is key to further investigate the underlying role of CAR cells in lung cancer spurred manipulation of hematopoiesis and involvement of S100A9 and LCN2.

## Conclusions

This study comprehensively profiled the BM niche of lung cancer bearing mice with validation studies on NSCLC patients. We thereby confirm that lung cancer can remotely alter the BM niche resulting in increased abundance of uni- and multi-potent progenitors at the expense of mature Neu and B cells. Furthermore, a significant increase in expression and secretion of S100A9 and LCN2 was observed in the BM of LLC-bearing mice, which we confirmed on the protein level in rib fragments of NSCLC patients. Finally, our data showed that S100A9 inhibition via Tasquinimod reduced lung tumor volumes in an aggressive orthotopic lung cancer model irrespective of anti-PD-L1 mAb treatment. As anti-PD-L1 mAb treatment further increased the levels of secreted S100A9 and LCN2 in the BM, Tasquinimod counteracted this effect for both. These findings highlight the potential of scrutinizing their potential role as biomarkers and therapy targets in lung cancer refractory to immunotherapy.

## Materials and Methods

### Cell line

Lewis lung carcinoma (LLC) cells, generously provided by Prof. Jo Van Ginderachter (VUB), were cultured in Dulbecco’s Modified Eagle’s Medium (DMEM, Gibco) supplemented with 10% foetal bovine serum (FBS, PAN Biotech), 62.5U/mL penicillin, 62.5µg/mL streptomycin, 2.5 mM L-glutamine (Sigma-Aldrich), 1.25 mM sodium pyruvate (ThermoFisher) and 1.25X MEM non-essential amino acids (Gibco). The supplemented medium is referred to as DMEM+. Cultures were maintained at 37°C in a humidified atmosphere containing 5% CO2, 21% O2 and 95% humidity.

### Mice and orthotopic LLC models

Six-week-old female and male C57BL/6J mice were purchased from Charles River. All experimental procedures, animal handling, and care were conducted in compliance with the Ethical Committee Dierproeven (ECD) from the Vrije Universiteit Brussel for use of laboratory animals (ECD file numbers: 20-214-12, 24-272-23, 24-272-35, 24-272-37, 25-272-8). To obtain flushed BM from orthotopic LLC-bearing mice for scRNA-seq analysis and humeri for immunohistochemistry (IHC), mice were i.v. injected with 200µL of PBS (Ctrl) or 5 × 10^5^ LLC cells. To obtain flushed BM for LSK sorting, LLC-bearing mice with localized orthotopic tumors were generated via intrathoracic challenge with 20 µL of 5×10^4^ LLC cells. For the survival experiment, mice were i.v. injected with 2,5 × 10^5^ LLC cells. Animal welfare was monitored twice per week by evaluation of weights and breathing rates. All animals were euthanized via cervical neck dislocation.

### Human samples

Rib fragments from operable NSCLC patients were collected at the Universitair Ziekenhuis Leuven as approved by the Medical Ethical Committee under the study protocol S68836.

### Tissue harvesting and processing

Immediately after animal euthanasia by cervical dislocation, blood was collected via cardiac puncture, animals were perfused with 10mL of PBS, and organs were subsequently harvested. *Lungs* were collected in a tube containing 3mL of DMEM+ medium with 30 U/mL of collagenase type I (Sigma-Aldrich). Lungs were chopped, digested at 37°C for 40 min, and mechanically reduced with an 18G needle. *Spleens* were collected in phosphate-buffered saline (PBS, Gibco) and gently dissociated by mechanical disruption. Murine femurs, tibiae, hips and spines were isolated and cleaned to remove soft tissue for BM collection. *Flushed BM* was isolated by excising both ends of the femurs and tibias, followed by centrifugation to obtain the BM. *Crushed BM* was obtained by grinding the hips, spine and flushed hind legs in digestion medium (RPMI 1640, Gibco) containing 60 U/mL of collagenase type IV(Gibco) and 0.5mg/mL of dispase II (Sigma-Aldrich) using a mortar and pestle as previously described (Baccin et al., 2019; S. Xu et al., 2010). Crushed bone chips with dissociated cells were incubated at 37°C for 30 min in digestion medium. Digestion was neutralized with 40 mL of RPMI 1640 medium. All cell suspensions were filtered through a 40 µm strainer, centrifuged (1500 rpm, 5 min), then incubated for 5 min with red blood cell lysis buffer. Neutralization was achieved by adding 5mL of PBS, centrifugation (1500 rpm, 5min) and resuspension in DMEM+. For RT-qPCR, 5 x 10^6^ crushed murine BM cells were snap-frozen and stored at - 80°C. Blood (100µL) was collected into 1.3 mL LH microtubes (Sarstedt) and centrifuged (2000g, 10 min) to separate cell pellets from plasma. Another 100 µL of blood was recovered into Eppendorf tubes containing heparin for subsequent RNA extraction.

### scRNA-seq – 10x genomics

Flushed BM were collected from C57BL/6J female mice, 3 weeks after their i.v. challenge with PBS or LLC (n=4). To eliminate dissociation-induced expression of immediate-early genes(Scheyltjens et al., 2022), all BM samples were processed in buffers containing Actinomycin D (ActD, Gibco) dissolved in sterile DMSO. Briefly, PBS with 30 µM ActD was used to collect bones, for further processing and washing steps PBS with 3 µM ActD was used. Finally, cells were resuspended in PBS with 0.04% BSA and 3µM ActD. Per mouse, 2 × 10^6^ BM cells were used to initiate barcoding using the 10X Genomics protocol for scRNA-seq. Cells from each mouse were tagged with cell multiplexing oligos (CMO). After labelling, 1×10^4^ cells per mouse were pooled together per condition (healthy and LLC). Reverse transcriptase PCR, library preparation and sequencing were performed via BrightCore-VUB.

### Analysis of scRNA-seq data

The Cell Ranger software (v.6.0.2, 10x Genomics) was used for sample demultiplexing, alignment of RNA reads to the mm10-2020A reference genome, processing of RNA and barcodes, filtering of unique molecular identifiers (UMI) and quantification of single-cell UMI counts. The RNA expression matrices were further filtered and processed using the Seurat (Satija et al., 2015) R package (v.4.0.3 and v.4.3.0). Empty droplets (contaminant ambient RNA) were identified using DropletUtils (Lun et al., 2019) package (v. 1.10.3) (niters = 5000), and droplets with false discovery rate (FDR) ≤ 0.01 were retained for downstream analysis. Datasets were normalized, and highly variable genes were identified, selecting the top 2000 variable genes per sample. Shared integration features were identified, and datasets were integrated using the anchor-based workflow (Stuart et al., 2019). The original RNA assay was retained for differential expression analyses. Cell-level quality control metrics included total UMI counts per cell, number of detected genes, and the percentage of mitochondrial gene expression. Outliers were identified using the median absolute deviation and cells with high mitochondrial content were excluded (McCarthy et al., 2017). Lowly expressed genes (mean < 0.005 UMIs across all samples) were removed. Doublets were identified using scDblFinder(Germain et al., 2022) package (v.1.4.0). Samples were demultiplexed and multiplets, blanks, and unassigned cells were excluded from downstream analyses (Stoeckius et al., 2018) Following filtering and demultiplexing, the integrated assay was scaled and principal component analysis (PCA) was performed. Uniform Manifold Approximation and Projection (UMAP) was performed using the first 30 PCs.

Subsequently, the identified highly variable genes were used for performing principal component analysis (PCA). The PCA embeddings were used downstream for unsupervised Leiden clustering of the cells and Uniform Manifold Approximation and Projection (UMAP) dimensionality reduction as implemented in Seurat. A shared nearest neighbor (SNN) graph was constructed using the first 30 PCs, and Leiden clustering (Traag et al., 2019) was applied. Resolution 0.6 was selected based on cluster stability and biological interpretability. Cluster-specific marker genes were identified using Seurat’s FindMarkers function on the non-integrated RNA assay. Cell annotation was supervised using canonical markers and CellKb(Patil & Patil, 2022). Differential gene expression was assessed per cluster between LLC-bearing and healthy samples. A minimum log₂ fold-change threshold of 0.8 and Bonferroni-adjusted P < 0.1 were used to define significant genes. Volcano plots were generated using EnhancedVolcano. Cell-type–specific transcriptional shifts were quantified using Cacoa (Petukhov et al., 2022) package, with healthy samples as the reference and LLC-bearing samples as the target. Intercellular communication was evaluated by NicheNet (Browaeys et al., 2019)ligand-receptor analysis while trajectory and pseudotime analysis was evaluated by Monocle3.

### Expression analysis of target genes by qPCR

RNA extraction from crushed murine BM cells was performed using the RNeasy Mini kit (Qiagen) according to the vendor’s instructions. In addition, five slices of 10µm from paraffinized human rib fragments were collected for RNA extraction using the PureLink FFPE Total RNA Isolation Kit. The quantity and purity of the extracted RNA were assessed using a NanoDrop (Thermo Fisher Scientific). cDNA conversion was performed with Verso cDNA Synthesis Kit (Thermo Scientific^TM^) according to the manufacturer’s instructions. PowerUP^TM^ SYBR^TM^ Green Master Mix for qPCR was used according to the manufacturer’s instructions, with the corresponding primers (Extended Data Table 2). For plate read-out, the QuantStudio 12K Flex Real-Time PCR System (Thermo Fisher Scientific) was used with the following protocol: Hold stage 94°C – 2 min, PCR stage 94°C – 15 sec, (40 cycles) 60°C – 1 min, Melt Curve 95°C – 15 sec, followed by 60°C – 1 min. Housekeeping genes used for normalization were *Atp5b* for mouse BM, *Rplp0* for mouse blood, and *GAPDH* for human BM samples.

### ELISA

Sera, lungs supernatants and BM supernatants were used to test secreted S100A9 (Biotechne, R&D systems), LCN2 (Thermo Fisher Scientific) and IL17A (Invitrogen), according to manufacturers instructions. Plates were read in the GloMax at 450nm.

### Flow cytometry

Cells were centrifuged (1500rpm, 5min) and washed with PBS. Then, Fc receptors were blocked with a purified anti-CD16/32 antibody (BioLegend), and cells were stained with ZombieAqua (BioLegend) following manufacturer’s instructions.. Cells were washed with FACS buffer (PBS containing 1% bovine serum albumin (BSA) and 0.02% sodium azide (Sigma-Aldrich)). Next, staining with an antibody cocktail was performed for 30 min at 4°C, in the dark, in FACS buffer. Cells were washed with FACS buffer and centrifuged (1500rpm, 5min). For intracellular staining, cells were fixed and permeabilized with BD Cytofix/Cytoperm^TM^ Fixation/Permeabilization Solution Kit (BD) following manufacturer’s protocol. After permeabilization cells were stained with intracellular cocktail in 1X permeabilization buffer (BD), for 30 min at 4°C in the dark. Fluorescently labelled cells were evaluated on a Symphony A1 flow cytometer (BD), and analysis was performed using Flowjo V10.10.0 software. The list and dilutions of antibodies can be found in Extended Data Table 3

### IHC and two-photon imaging of fixed bone

Murine humeri were isolated, cleaned, washed in PBS and placed in 4% paraformaldehyde at 4°C overnight. Subsequently, the fixed humeri were washed in PBS and decalcified in 0.45 M EDTA, pH 7.4, at 4°C for 5 days. Dehydration was achieved by overnight incubation at 4°C in 15% sucrose, followed by 1 day in 30% sucrose. Humeri were washed in PBS, embedded in PolyFreeze Tissue Freezing medium (Sigma-Aldrich), and stored at -80 °C until cryotome sectioning. Sections of 25 µm were transferred to Epredia^TM^ SuperFrost Plus^TM^ adhesion slides.

Human rib fragments were collected from operable patients at Universitair Ziekenhuis Leuven (UZL). Rib fragments were fixed in 4% paraformaldehyde at 4°C overnight. Subsequently, fixed ribs were washed in PBS and decalcified in formic acid for 5 days, processed by the histokinette and embedded in paraffin for 2 h. Paraffin blocks were soaked in cold 0.45 M EDTA, pH7.4, before microtome sectioning at 25 µm and transfer to Superfrost adhesion slides. Samples were deparaffinized by heating in an oven at 60°C for 1 h, followed by washings in 2 × toluene, 2 × ethanol 100%, 1 × ethanol 90%, and 1 × ethanol 70%. Antigen retrieval was performed with citrate buffer (ImTec) in a steamer for 25 min.

Sections were washed once for 5 min at room temperature (RT) in PBS containing 0.1% Tween 20 (PBST), then blocked for 2 hours at RT with blocking buffer (PBS with 10% FBS and 2% BSA). Slides were washed 3 x in PBST at RT, 5 min each. Then, samples were stained overnight at 4°C with antibodies in PBST with 10% blocking buffer. Following incubation, slides were washed three times for 5 min at RT with PBST, incubated for 2 hours at RT with secondary antibodies diluted in PBST 10% blocking buffer, and then washed three times in PBST. Finally, samples were mounted using Fluoroshield^TM^ histology mounting medium with DAPI. The list of antibodies and their corresponding dilutions can be found in Extended Data Table 3.

Fixed bones were imaged using a Stellaris 8 DIVE two-photon microscope (Leica Microsystems) with an upright objective at 25X magnification, acquired with the LAS X Life Science Microscope Software Platform. Microscopy images were analyzed using Fiji.

### Intravital two-photon imaging of BM calvarium

Flushed BM cells were obtained from C57BL/6J female mice, 3 weeks after their intrathoracic challenge with PBS or LLC cells. A total of 6 × 10^4^ LSK cells per mouse were sorted (BD FACSAria^TM^ III) and labelled with cell trace violet (CTV, Life Technologies) prior to their i.v. injection into healthy C57BL/6J mice. Seven days later, mice were i.v. injected with 5µg of Brilliant Violet 711 anti-mouse cKit antibody (BioLegend, clone: 2B8).

For *in vivo* imaging, the protocol was adapted from (J. W. Wu et al., 2014). Briefly, mice received subcutaneously 50 µL of Vetergesic (buprenorphine 0.05 mg/kg, diluted 7.46x) and intraperitoneally 500 µL of 0.9% NaCl 30 min before surgery. Mice were anesthetized with isoflurane (5% induction, 2,5% maintenance, oxygen flow rate between 0.3 and 1.5 L/min). Eye ointment was applied, followed by shaving of whiskers, head and forehead, including the areas adjacent to the eyes. Depilatory cream was applied for up to 2 min to ensure hair removal. Xylocaine gel was applied for topical analgesia, 20 min before the skin was disinfected with iso-Betadine and chlorhexidine. A triangular incision was made on the forehead skin using surgical scissors, with the vertex positioned near the nasal bridge. The first cut extended laterally near the eye to form one side of the triangle, followed by a second cut along the forehead to form the base. The resulting skin flap was gently retracted and adhered to the eye ointment. The exposed skull was kept moist with 0.9% NaCl, and the periosteal membrane was carefully removed using fine tweezers. The skull surface was cleaned for approximately 5 minutes to remove all residual tissue, ensuring optimal imaging conditions. A coverslip with stabilizing ring was placed over the cleaned and exposed skull. The mouse was fixed using a SGM-4 head holder (Narishige) and placed on a heating pad. Isoflurane-based anesthesia was maintained during imaging. To visualize vasculature, mice were i.v. injected with 30mg/kg 500kD FITC-Dextran (Invitrogen). Thirty minutes later, calvaria were imaged using a Stellaris 8 DIVE two-photon microscope (Leica Microsystems) with upright objective at 25X magnification, acquired with the LAS X Life Science Microscope Software Platform. Microscopy images were analyzed using Fiji. CTV and c-Kit signals were quantified by 3D Objects counter tool.

### *In vivo* survival experiment

An equal amount of male and female 6-week-old mice were i.v. challenged with 2.5 × 10^5^ LLC cells (n=8). Tumor volumes were monitored every 3-4 days via micro-CT imaging (Molecubes). Seven days after LLC challenge, mice were divided in 4 groups: Isotype Control (IC), anti-PD-L1 mAb (IT), IC with Tasquinimod and IT with Tasquinimod. In brief, mice were injected intraperitoneally for 4 times every 3-4 days with 200 µg/ 200 µl PBS of Ultra-LEAF™ Purified anti-mouse CD274 (PD-L1) rat IgG2b, K mAb (BioLegend, clone: 10F.9G2) or Ultra-LEAF™ Purified rat IgG2b, K isotype control (Ctrl) mAb (BioLegend, clone: RTK4530). For Tasquinimod (MedChemExpress) treated mice, 30mg/kg was administered via drinking water from day 7 after tumor challenge. Progression of the disease was monitored by micro-computer tomography (micro-CT), body weight and breathing rate three times per week until the humane endpoint was reached.

### micro-CT imaging

Mice were sedated with isoflurane (5% induction, 2,5% maintenance, oxygen flow rate between 0.3 and 1.5 L/min) and placed on a bed for micro-CT imaging (Molecubes, 200 mGy for 5min). CT data were processed for tumor manual segmentation and volume estimation via 3D slicer (Kikinis et al., 2014).

### Statical analysis

Analysis of micro-CT images were performed blindly. For the rest of the experiments, data collection and analysis were not performed blindly to the conditions of the experiments. For experiments involving tumor induction, female and male C57BL/6J mice were randomly assigned to control, disease or treated group. Statistical tests are described in the legends of each figure. In the absence of statistical significance, no *P* value was plotted on the figures. Statistical analysis of the scRNA-seq data were performed using R Studio (4.3.0), while all other statistical analyses were conducted using GraphPad Prism v10.

## Acknowledgements

E.C.E’s work is supported by the EUTOPIA PhD Co-tutelle Programme. This research was further performed with the financial support of Wetenschappelijk Fonds Willy Gepts of the UZ Brussel and Research Foundation-Flanders (FWO-V, grant I002422N and G041721N).

The authors acknowledge the *In vivo* Cellular and Molecular Imaging (ICMI) Core Facility at Vrije Universiteit Brussel (VUB) for providing access to the *in vivo* preclinical imaging infrastructure, in particular the two-photon intravital microscope acquired via FWO grant I002422N. The authors thank Sofie Pollenus, Maxime Deladriere and Kevin De Jonghe for their technical assistance.

The authors also acknowledge the Flow Cytometry Core Facility (FlowCore) and Visual & Spatial Tissue Analysis (VSTA) Core Facility at VUB for their support in flow cytometry and tissue sectioning. The authors thank Emmy De Blay for her technical assistance.

## Author contributions

All authors researched data for the article, contributed to discussions on its content and drafted the manuscript under the guidance of C.G. All authors reviewed and approved the final version before submission.

