## Extended Data Fig. for "Lung cancer-fueled emergency myelopoiesis is characterized by an increase of S100A9^+^ and LCN2^+^ hematopoietic stem and progenitor cells"

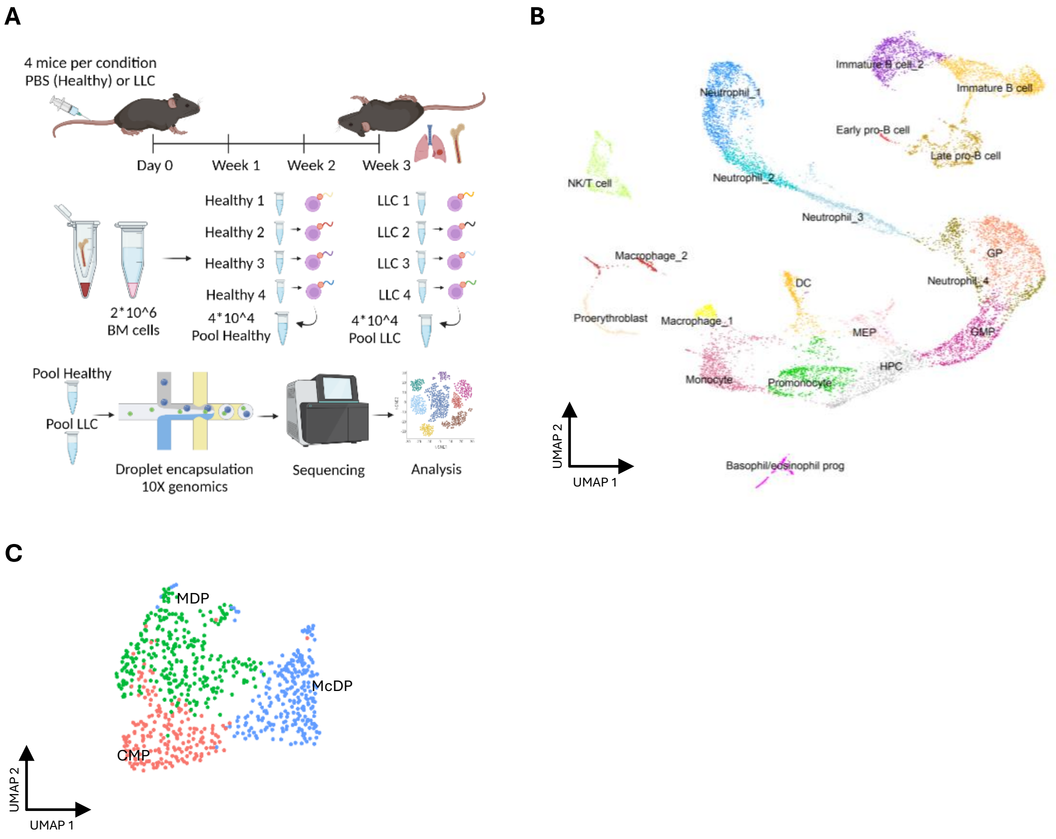


**Extended Data Fig. 1 Hashed scRNA-seq experiment. (A)** Setup of the hashed scRNA-seq experiment. scRNA-seq analysis was performed on whole BM isolated from hind legs of healthy female (n=4) and LLC-bearing female mice (n=4). **(B)** Hashed scRNA-seq data from healthy and LLC-bearing mice were processed and integrated using Seurat's (v.4.3.0) anchor-based integration approach. Harmony-corrected UMAP plots of 6674 and 6867 BM cells resp. showed 21 clusters at 0.6 resolution. **(C)** Annotations of subclusters when zoomed in on HP-related cell type at 0.3 resolution. Panel **A** created with BioRender.com.

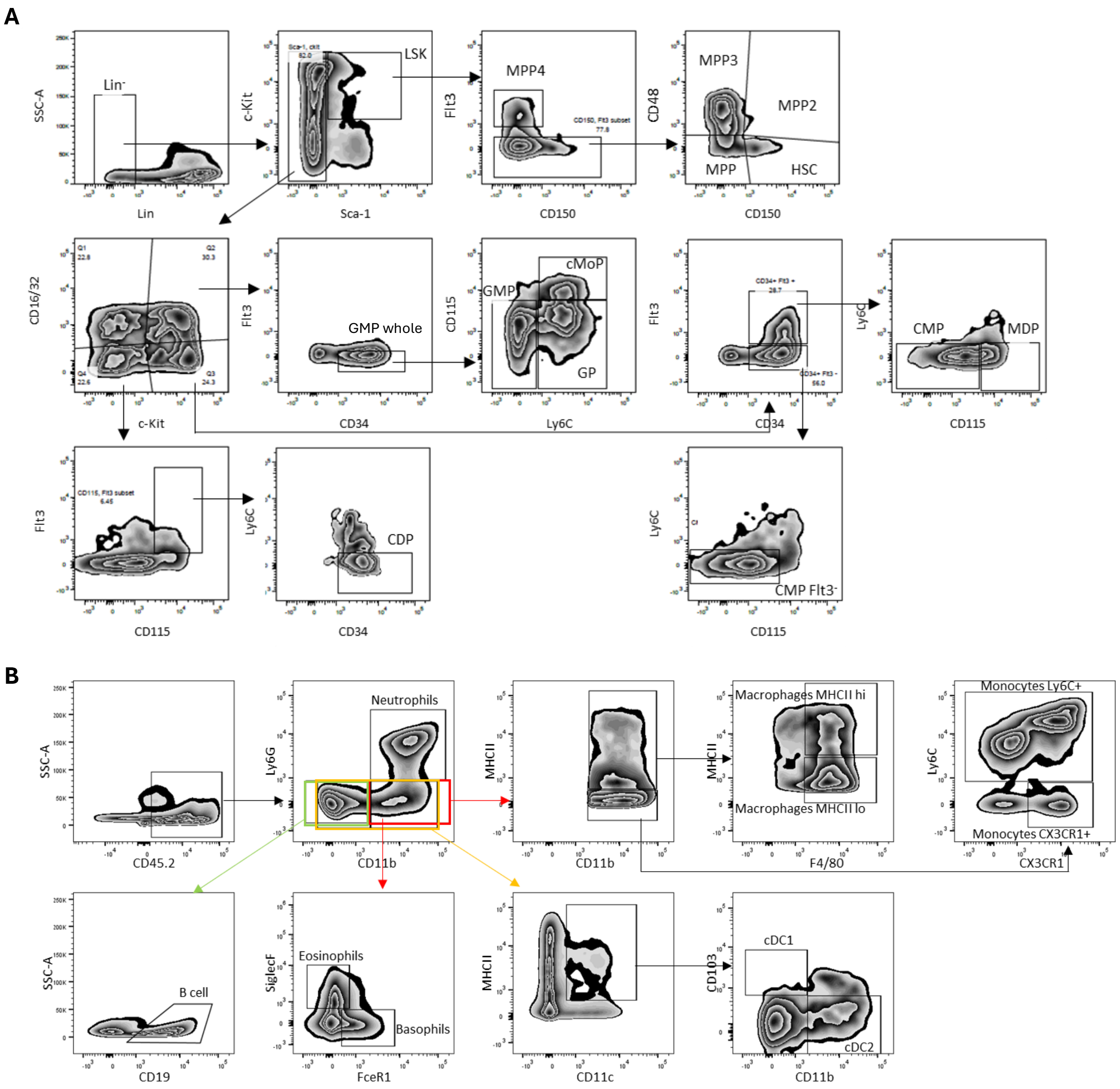


**Extended Data Fig. 2 Gating strategies to characterize HSPC and differentiated immune cell subsets via flow cytometry. (A)** Lin$^-$ cells were stained for Sca-1, c-Kit, CD150, Flt3, CD48, CD16/32, CD34, CD115, Ly6C, CD45 and CD105 to distinguish: Multipotent Progenitor (MPP)-Lymphocyte/Granulocyte/Megakaryocyte-erythroid (MPP4/MPP3/MPP2), Hematopoietic Stem Cells (HSC), Granulocyte Monocyte Progenitor (GMP), Granulocyte Progenitor (GP), Common Monocyte Progenitor (cMoP), Monocyte-DC Progenitor (MDP), Common Myeloid Progenitors (CMP), Common Dendritic cell Progenitor (CDP) and stromal cells. **(B)** To identify neutrophils, basophils, eosinophils, macrophages, dendritic cells, monocyte and B cells: CD45+ cells were stained for Ly6G, CD11b, MHC II, F4/80, Ly6C, CX3CR1, CD11c, CD103, Siglec F, FceR1 and CD19.

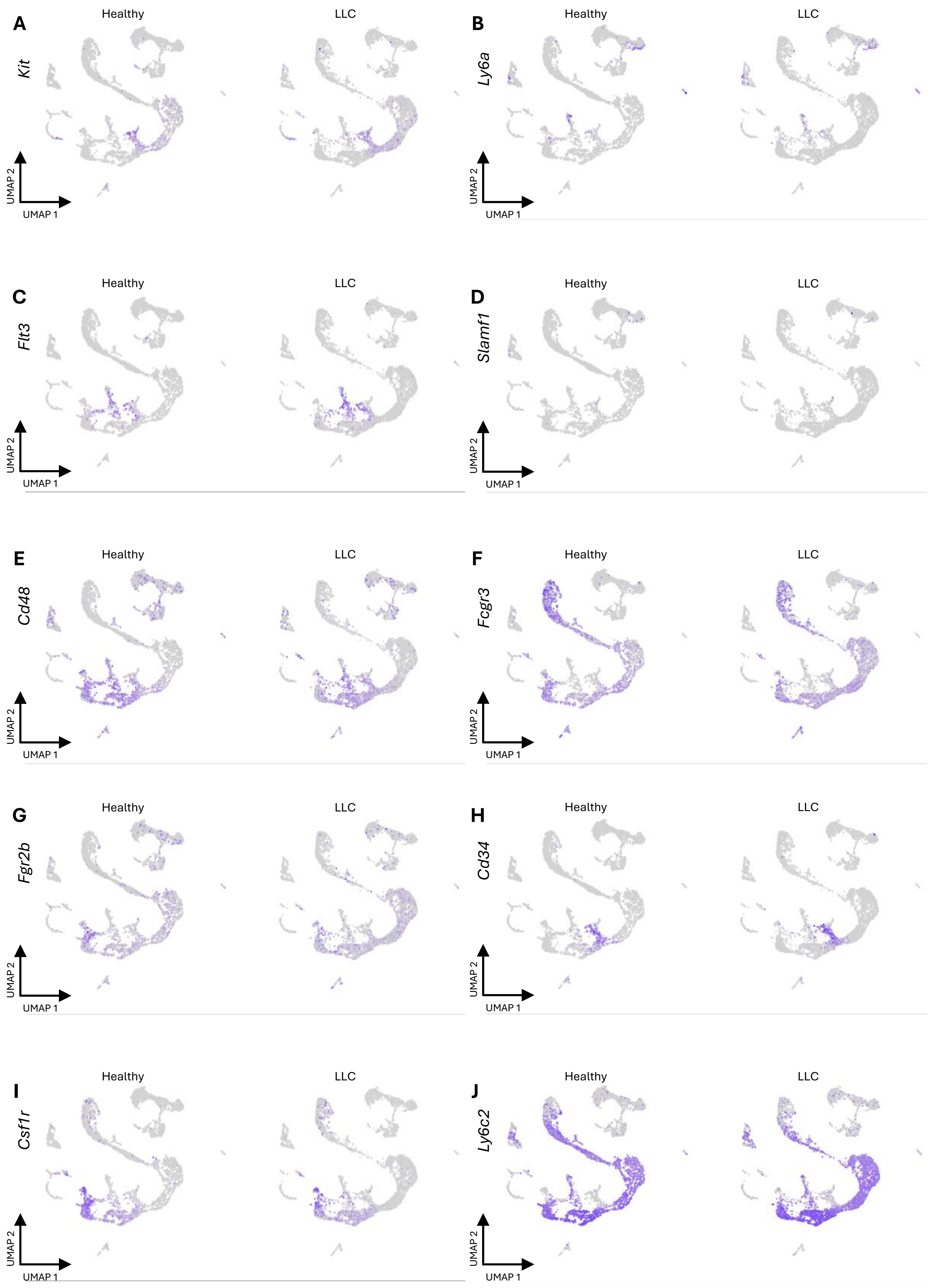


**Extended Data Fig. 3 Feature plots depicting the expression of flow cytometry markers in HSPC panel across individual cells in healthy and LLC-bearing mice**. Blue color intensity represents the expression level. **(A)** *Kit* corresponds to c-Kit. **(B)** *Ly6a* corresponds to Sca-1. **(C)** *Flt3* corresponds to Flt3. **(D)** *Slamf1* corresponds to CD150. **(E)** *Cd48* corresponds to CD48. **(F)** *Fcgr3* corresponds to CD16. **(G)** *Fcgr2b* corresponds to CD32. **(H)** *Cd34* corresponds to CD34. **(I)** *Csf1r* corresponds to CD115. **(J)** *Ly6c2* corresponds to Ly6C.

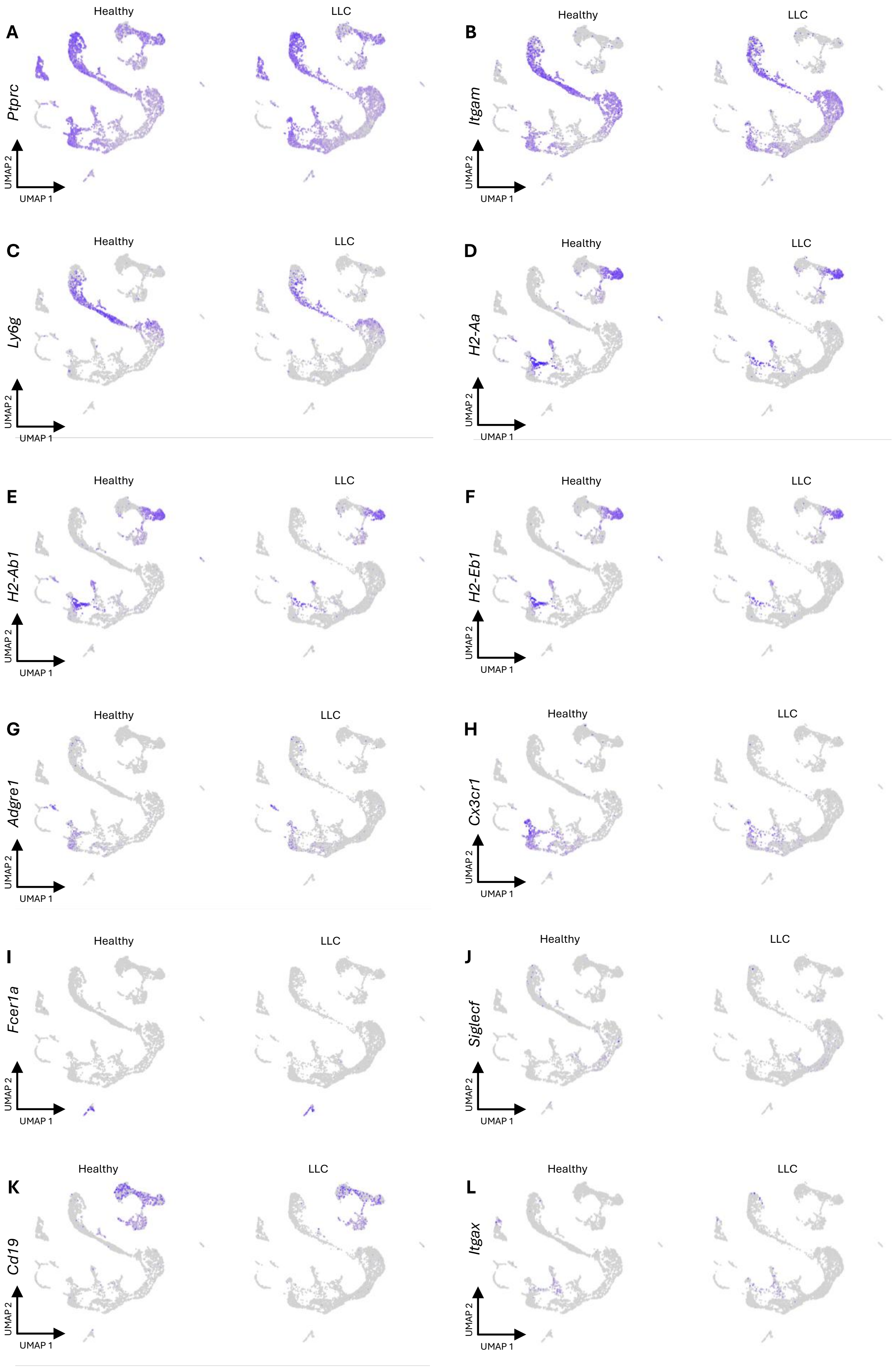
A
Ptprc
B
Itgam
C
Ly6g
D
H2-Aa
E
H2-Ab1
F
H2-Eb1
G
Adgre1
H
Cx3cr1
I
Fcer1a
J
Siglecf
K
Cd19
L
Itgax
Healthy
LLC
UMAP 1
UMAP 2

**Extended Data Fig. 4 Feature plots depicting the expression of flow cytometry markers in differentiated immune cells panel across individual cells in healthy and LLC-bearing mice**. Blue color intensity represents the expression level. **(A)** *Ptprc* corresponds to CD45. **(B)** *Itgam* corresponds to Cd11b. **(C)** *Ly6g* corresponds to Ly6G. **(D)** *H2-Aa,* **(E)** *H2-Ab1,* and **(F)** *H2-Eb1* corresponds to MHCII. **(G)** *Adgre1* corresponds to F4/80. **(H)** *Cx3cr1* corresponds to CX3CR1. **(I)** *Fcer1a* corresponds to FceR1. **(J)** *SiglecF* corresponds to Siglec F. **(K)** *Cd19* corresponds to CD19. **(L)** *Itgax* corresponds to CD11c.

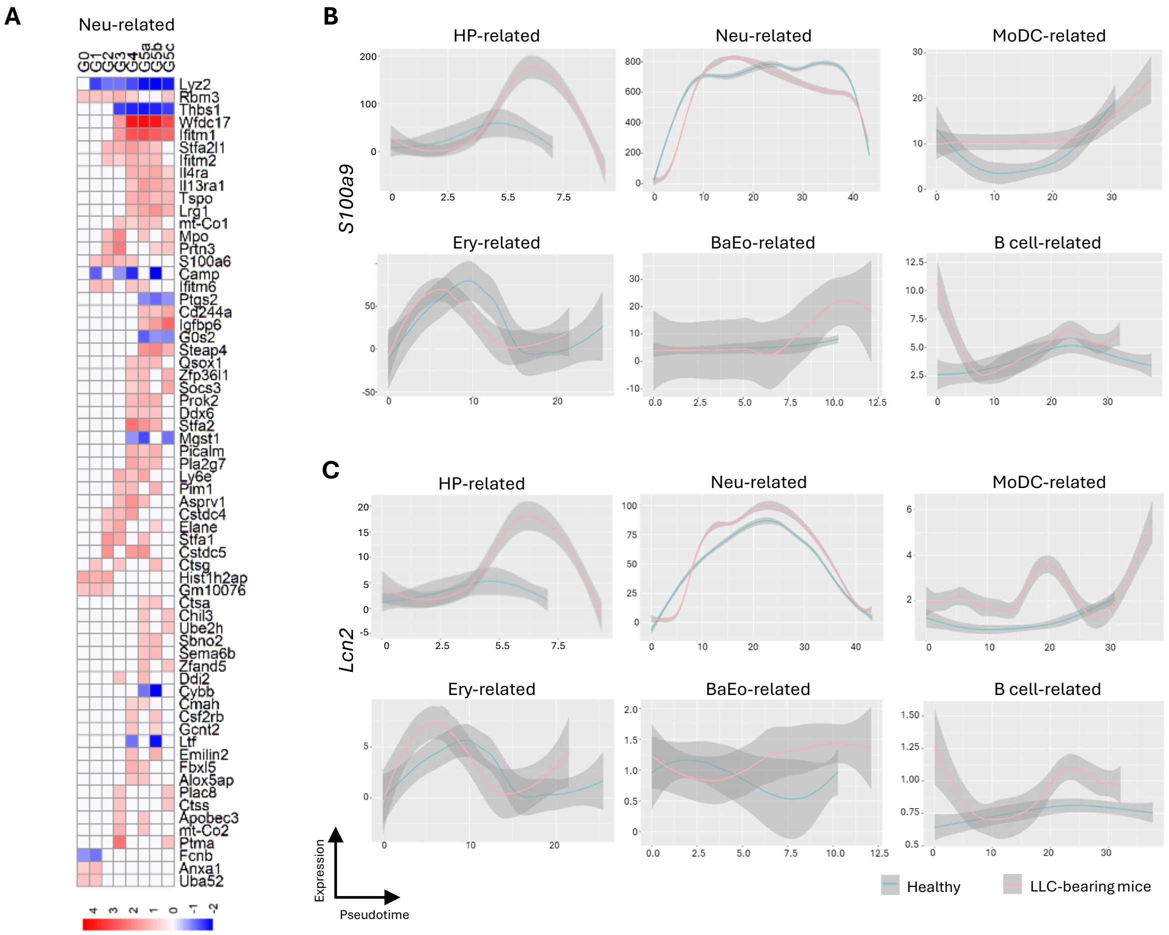


**Extended Data Fig. 5** ***S100a9*** **and** ***Lcn2*** **are upregulated in all subclusters except for the Neu-related cluster. (A)** Heatmap of ranked up- and down-regulated genes, significant in 2 or more subsets within the Neu-related cluster. Adjusted p-value< 0.1 and |Log2FC| > 0.8, red for up- and blue for down-regulated genes in LLC-bearing versus healthy mice. Plots show the expression of **(B)** *S100a9* and **(C)** *Lcn2* as a function of pseudotime along the hematopoietic trajectory in the whole data set and specific cell types. Lines show smoothed expression trends, and shaded areas indicate confidence intervals.

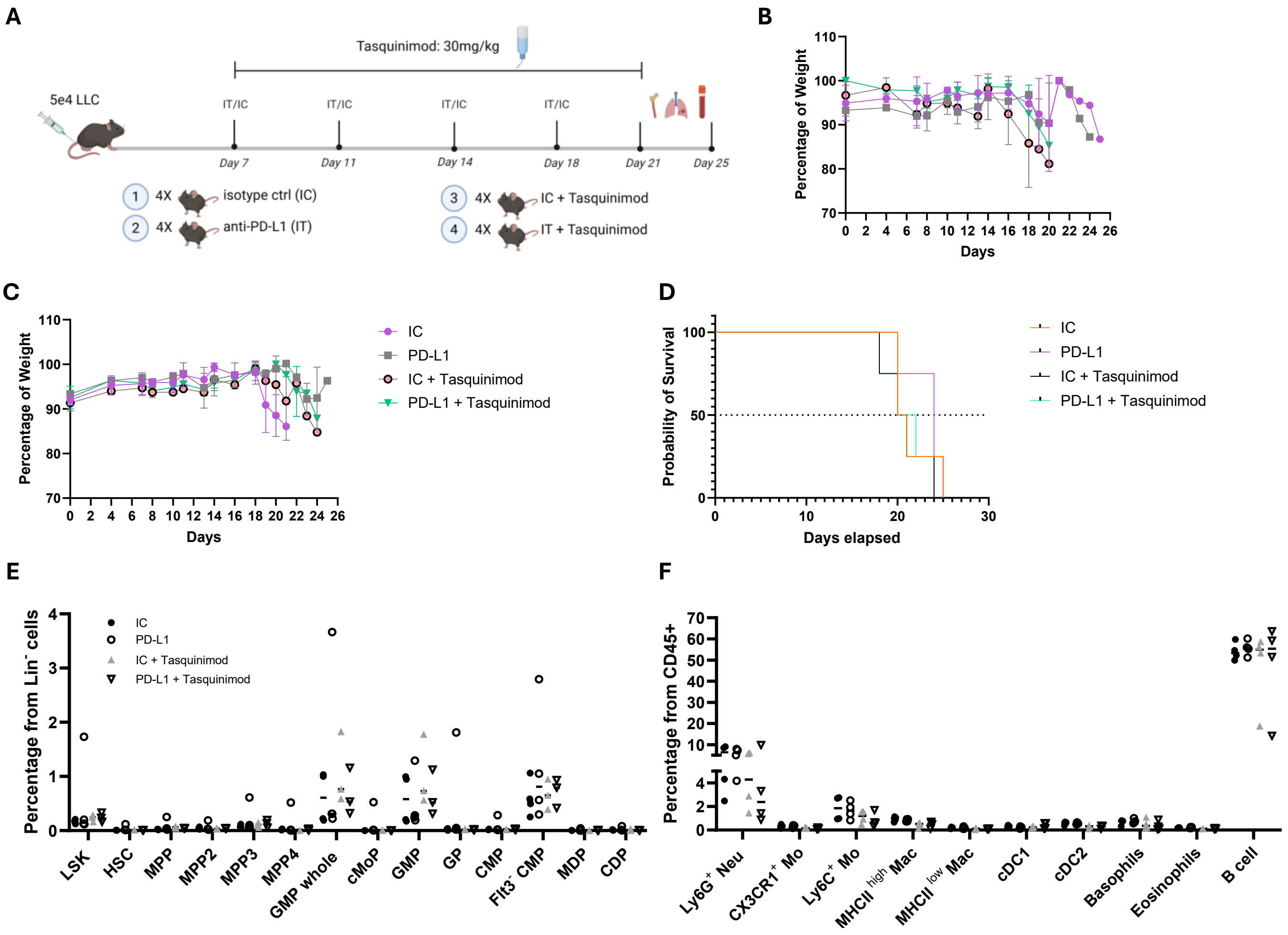


**Extended Data Fig. 6 Welfare monitoring during the Tasquinimod *in vivo* experiment. (A)** Experimental setup of the gender-balance *in vivo* Tasquinimod experiment. Mice were i.v. challenged with 5x10[4] LLC cells. LLC-bearing mice were treated with IC (n=2F, 2M), anti-PD-L1 mAb (n=2F, 2M), IC + Tasquinimod (n=2F, 2M) or anti-PD-L1 mAb + Tasquinimod (n=2F, 2M). **(B-C)** Longitudinal body weight measurements over a period of 25 days in **(B)** female and **(C)** male mice. **(D)** Kaplan Meier survival curve for all animals under the 4 different treatments. Dot plots depict *ex vivo* spleen flow cytometry abundances of **(E)** Lin⁻ progenitor subsets and **(F)** CD45⁺ differentiated immune cells. Depicted p-values were obtained upon Mann-Whitney U test.

**Extended Data Table 1. Mann-Whitney U test p-values**

| **Cell type** | ***P* value** | **q value** |
|---|---|---|
| HSC | 0.21927 | 0.221462 |
| MPP | 0.002176 | 0.007325 |
| MPP2 | 0.001554 | 0.007325 |
| MPP3 | 0.013986 | 0.023543 |
| MPP4 | 0.054079 | 0.078029 |
| GMP whole | 0.005905 | 0.014911 |
| cMoP | >0.999999 | 0.776923 |
| GMP | 0.000311 | 0.003139 |
| GP | 0.009324 | 0.018834 |
| CMP | >0.999999 | 0.776923 |
| Flt3- CMP | 0.072106 | 0.080919 |
| MDP | 0.713287 | 0.654927 |
| CDP | 0.072106 | 0.080919 |
| Ly6G$^{+}$ Neu | 0.040093 | 0.122895 |
| CX3CR1$^{+}$ Mo | 0.463403 | 0.702056 |
| Ly6C$^{+}$ Mo | 0.778866 | 0.786654 |
| MHCII $^{high}$ Mac | 0.054079 | 0.122895 |
| MHCII $^{low}$ Mac | 0.630769 | 0.786654 |
| cDC1 | 0.093862 | 0.170641 |
| cDC2 | 0.054079 | 0.122895 |
| Basophils | 0.866511 | 0.787659 |
| Eosinophils | 0.694328 | 0.786654 |
| B cell | 0.000311 | 0.002825 |

**Extended Data Table 2. List of antibodies**

| Antigen | Specificity | Conjugation | Clone | Dilution | Source | App |
|---|---|---|---|---|---|---|
| Lcn2 (Ngal) | Mouse | AF488 | H-7 | 50 | Santa Cruz | FC/IF |
| S100a9 | Mouse | AF647 | 2B10 | 50 | BD Pharmigen | FC/IF |
| Ly6C | Mouse | AF594 | HK1.4 | 50 | Biolegend | FC/IF |
| c-Kit | Mouse | unconjugated | polyclonal | 50 | Thermo Fisher Scientific | IF |
| Rabbit IgG | Rabbit | Cy3 | | 400 | Jackson Immunoresearch | IF |
| CD34 | Human | AF488 | 561 | 50 | Biolegend | FC/IF |
| S100A9 | Human | PE | MRP 1H9 | 50 | Biolegend | FC/IF |
| LCN2 (NGAL) | Human | unconjugated | polyclonal | 200 | Proteintech | IF |
| Rabbit IgG | Rabbit | AF647 | | 400 | Jackson Immunoresearch | IF |
| CD34 | Mouse | BV421 | SA376A4 | 200 | Biolegend | FC |
| CD16/32 | Mouse | FITC | 93 | 200 | Biolegend | FC |
| Flt3 (CD135) | Mouse | PE | A2F10 | 200 | Biolegend | FC |
| CD115 | Mouse | BV605 | AFS98 | 200 | Biolegend | FC |
| CD11c | Mouse | PE-Dazzle | N418 | 200 | Biolegend | FC |
| CD19 | Mouse | PE-Dazzle | 6D5 | 200 | Biolegend | FC |
| CD3e | Mouse | PE-Dazzle | 145-2C11 | 200 | Biolegend | FC |
| CD49b | Mouse | PE-Dazzle | DX5 | 200 | Biolegend | FC |
| Ly6G | Mouse | PE-Dazzle | 1A8 | 200 | Biolegend | FC |
| Ter-119 | Mouse | PE-Dazzle | TER-119 | 200 | Biolegend | FC |
| CD11b | Mouse | PE-Dazzle | M1/70 | 200 | Biolegend | FC |
| CD45R/B220 | Mouse | PE-Dazzle | RA3-6B2 | 200 | Biolegend | FC |
| Ter119 | Mouse | PerCPCy5.5 | TER-119 | 200 | Biolegend | FC |
| SCA-1 (Ly6A/E) | Mouse | AF700 | D7 | 200 | Biolegend | FC |
| CD335 (NKp46) | Mouse | BV605 | 29A1.4 | 200 | Biolegend | FC |
| CD209a (DC-SIGN) | Mouse | PE | MMD3 | 200 | Biolegend | FC |
| CD150 | Mouse | BV650 | TC15-12Ff12,2 | 200 | Biolegend | FC |
| c-kit (CD117) | Mouse | BV711 | 2B8 | 200 | Biolegend | FC |
| CD105 | Mouse | BV785 | MJ7/18 | 200 | Biolegend | FC |
| Sca1 | Mouse | PE-Cy7 | D7 | 200 | Biolegend | FC |
| CD45.2 | Mouse | AF700 | 104 | 100 | Biolegend | FC |
| CD48 | Mouse | APC-Fire750 | HM48-1 | 200 | Biolegend | FC |
| CD19 | Mouse | BV650 | 6D5 | 200 | Biolegend | FC |
| Ly6G | Mouse | BV711 | 1A8 | 400 | Biolegend | FC |
| CD11b | Mouse | BV785 | M1/70 | 200 | Biolegend | FC |
| CD11c | Mouse | PE-Dazzle594 | N418 | 200 | Biolegend | FC |
| Ly6C | Mouse | APC-Fire750 | HK1.4 | 200 | Biolegend | FC |
| MHCII | Mouse | FITC | M5/114.15.2 | 200 | Biolegend | FC |

**Extended Data Table 3. Primer sequences**

| Specie | Gene | Primer sense | Sequence 5' → 3' |
|---|---|---|---|
| Mouse | *S100a9* | Forward | GCAAGAAGATGGCCAACAAAG |
| | | Reverse | GGTGTCCTTCCTTCCTAGAGTA |
| | *Lcn2* | Forward | TTTGTTCCAAGCTCCAGGGC |
| | | Reverse | CGTTCCTTCAGTTCAGGGGA |
| | *Il17a* | Forward | CAGAAGGCCCTCAGACTACC |
| | | Reverse | TCTCGACCCTGAAAGTGAAG |
| | *Atp5b* | Forward | GCCCGAGGAGTGCAAAAG |
| | | Reverse | CACGGCTTCTTCAATGGGTC |
| | *Rplp0* | Forward | GTCCTCGTTGGAGTGACATCG |
| | | Reverse | GTCTGCTCCCACAATGAAGC |
| | *Cxcl12* | Forward | CCTTCAGATTGTTGCACGGC |
| | | Reverse | CTTGCATCTCCCACGGATGT |
| Human | *S100A9* | Forward | GGAATTCAAAGAGCTGGTGC |
| | | Reverse | TCAGCATGATGAACTCCTCG |
| | *LCN2* | Forward | CCAATCGGTAATGGCCAGTCT |
| | | Reverse | CCTGTTTACCAGTCCAGGCA |
| | *IL17A* | Forward | CCCGGACTGTGATGGTCAAC |
| | | Reverse | GCTCTTCACAGTGGTCCTTCC |
| | *GAPDH* | Forward | GTGGCTGGCTCAGAAAAAGG |
| | | Reverse | GGGGAGATTCAGTGTGGTGG |
| | *CXCL12* | Forward | TGCCCTTCAGATTGTAGCCC |
| | | Reverse | CGAGTGGGTCTAGCGGAAAG |
